# Host-associated microbe PCR (hamPCR): accessing new biology through convenient measurement of both microbial load and community composition

**DOI:** 10.1101/2020.05.19.103937

**Authors:** Derek S. Lundberg, Pratchaya Pramoj Na Ayutthaya, Annett Strauß, Gautam Shirsekar, Wen-Sui Lo, Thomas Lahaye, Detlef Weigel

**Affiliations:** Department of Molecular Biology, Max Planck Institute for Developmental Biology, 72076 Tübingen, Germany; Department of Evolutionary Biology, Max Planck Institute for Developmental Biology, 72076 Tübingen, Germany; ZMBP-General Genetics, University of Tübingen, Auf der Morgenstelle 32, 72076 Tübingen, Germany

**Author notes:** Corresponding authors (D.W.) and (D.L.).

## Abstract

The ratio of microbial population size relative to the amount of host tissue, or “microbial load”, is a fundamental metric of colonization and infection, but it cannot be directly deduced from microbial amplicon data such as 16S rRNA gene counts. Because conventional methods to determine load, such as serial dilution plating or quantitative PCR, add substantial experimental burden, they are only rarely paired with amplicon sequencing. Alternatively, whole metagenome sequencing of DNA contributed by host and microbes both reveals microbial community composition and enables determination of microbial load, but host DNA typically greatly outweighs microbial DNA, severely limiting the cost-effectiveness and scalability of this approach. We introduce host-associated microbe PCR (hamPCR), a robust amplicon sequencing strategy to quantify microbial load and describe interkingdom microbial community composition in a single, cost-effective library. We demonstrate its accuracy and flexibility across multiple host and microbe systems, including nematodes and major crops. We further present a technique that can be used, prior to sequencing, to optimize the host representation in a batch of libraries without loss of information. Because of its simplicity, and the fact that it provides an experimental solution to the well-known statistical challenges provided by compositional data, hamPCR will become a transformative approach throughout culture-independent microbiology.

## Introduction

Knowing the relative abundance of individual taxa reveals important information about any ecological community, including microbial communities. An expedient means of learning their composition in a sample is to sequence and count a defined number of 16S or 18S rRNA genes (hereafter rDNA), the internal transcribed spacer (ITS) of rRNA arrays, or other amplicons that distinguish microbial species in a sample. However, these common amplicon counting-by-sequencing methods do not provide information on the density or load of the microbes. Critically, such microbial sequence counts lack a denominator accounting for the amount of the habitat sampled, and thus, sparsely-colonized and densely-colonized samples become indistinguishable, despite most study systems being open systems in which the total number of microbial cells can vary over many orders of magnitude. Another limitation of such compositional data is that because the sum of all microbes is constrained, an increase in the abundance of one microbe reduces the relative abundance of all other microbes, creating misleading interpretations in the absence of appropriate statistical methods^1–4^. Experimental determination of the microbial load, for example by relating microbial abundance to sample volume, mass, or surface area, has led to important insights in microbiome research that otherwise would have been missed with relative abundance data^5–12^.

For many host-associated microbiome samples, in particular those from plants^12^, nematodes^13^, insects^14–16^, and other organisms in which it is difficult or impossible to physically separate microbes from host tissues, a thorough DNA extraction yields both host and microbial DNA. For such samples, the amount of DNA from host and microbe is directly proportional to the number of cells sampled^17,18^, and therefore the ratio of microbial DNA to host DNA is an intrinsic measure of the microbial load of the sample^6,12,19–21^. Researchers have attempted to exploit this property and use the host rDNA amplified as a byproduct of microbial rDNA to calculate microbial load^6,21^, but because host nuclear ribosomal arrays may have hundreds or thousands of copies^22^, and organellar DNA is also overabundant, these methods are inefficient and require noisy interventions to increase the microbial signal. Sufficiently deep whole metagenome sequencing (WMS) also can in principle describe the microbial community composition and measure the microbial load, but is rarely practical because of a similar overrepresentation of host DNA^12,19^. For example, WMS of a leaf extract from wild *Arabidopsis thaliana* typically yields >95% plant DNA and <5% microbial DNA. Furthermore, many WMS reads remain unclassifiable and thus unquantifiable in complex samples^12,19^.

Most commonly, researchers combine amplicon sequencing with an additional orthogonal method. These include supplementary shallow WMS ^12^, quantitative PCR (qPCR) or digital PCR of host and/or microbial genes^3,16,19,23–26^, adding sequenceable “spike-ins” calibrated based on sample volume^27^, mass^10^, or qPCR-determined host DNA content ^25^, counting colony forming units (CFU)^11,28^, and flow cytometry^5,7^. The multitude of methods and publications hints at the enduring nature of this problem. While combining amplicon sequencing with any of these other approaches improves data, it requires more work, consumes more sample material, and introduces technical caveats, such as a reliance on accurately pipetting small quantities.

Here, we introduce host-associated microbe PCR or “hamPCR”, a robust and accurate single-reaction method to co-amplify a low-copy host gene and one or more microbial regions, such as 16S rDNA. We accomplish this with a two-step PCR protocol^29–33^. In hamPCR, gene-specific primer pairs bind to the ‘raw’ templates in a first short step, which is run for only two cycles to limit propagating amplification biases related to primer annealing and primer availability. In the second exponential step, a single set of primers with complementarity to the universal overhangs add barcodes and sequencing adapters. Such co-amplification of diverse fragments is used in many RNA-seq and WMS protocols^34,35^. Notably, Carlson and colleagues^29^ similarly used a two-step PCR including a multiplexed first step of five to seven cycles to sequence and quantify both variable and joining segments at human T and B cell receptor loci, providing strong proof-of-concept for our method applied to the microbiome.

We designed our host and microbe amplicons to have slightly different lengths, such that they can be resolved by electrophoresis for quality control. We further show that after pooling finished sequencing libraries, the amplicons can be separately purified and re-mixed at any favorable ratio prior for sequencing (for example, with host DNA representing an affordable 5-10%), and sequence counts can be accurately scaled back to original levels *in-silico*. Thus, in stark contrast to shotgun sequencing, samples with initially unfavorable host-to-microbe ratios can be easily adjusted *prior to sequencing* without loss of information. Because of the practical simplicity and flexibility of hamPCR, it has the potential to supplant traditional microbial amplicon sequencing in host-associated microbiomes.

## Results

### hamPCR generates quantitative sequencing-based microbial load

The first 2-cycle “tagging” step of hamPCR multiplexes two or more primer pairs in the same reaction, at least one of which targets a single- or low-copy host gene (Supplementary Figure 1). The tagging primers are then cleaned with Solid Phase Reversible Immobilisation (SPRI) magnetic beads^36^ (Supplementary Figure 2). Next, an exponential PCR of 20-30 cycles is performed using universal barcoded primers (Figure 1a, Supplementary Figure 1). As a host amplicon in *A. thaliana* samples, we targeted a fragment of the *GIGANTEA* (*GI*) gene, which is well conserved and present as a single copy in *A. thaliana* and many other plant species^37^. As microbial amplicons, we initially targeted widely-used regions of 16S rDNA.

**Figure 1.**
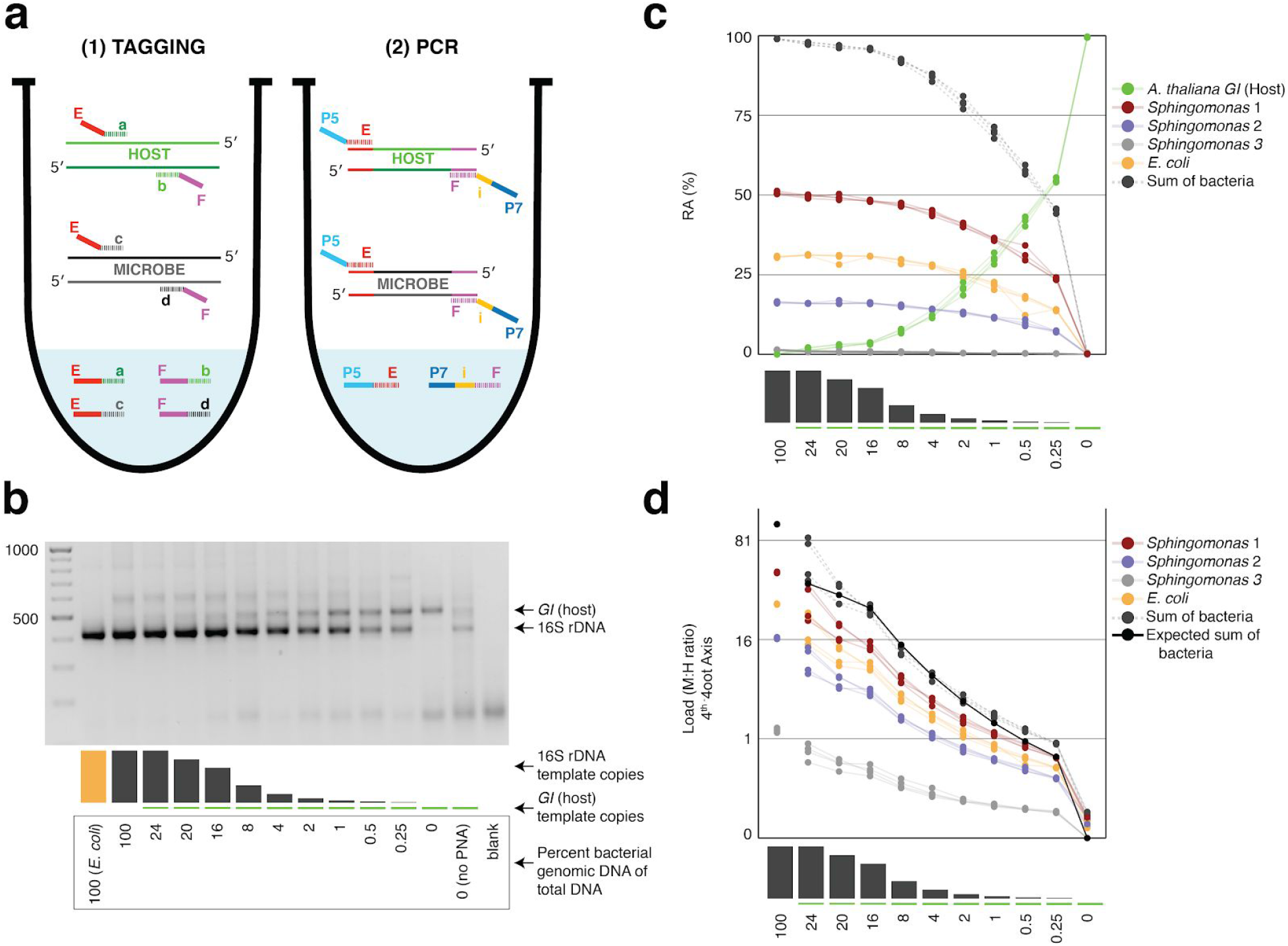
Synthetic samples demonstrate technical reproducibility. **a**, Schematic showing the two steps of hamPCR. The tagging reaction (left) shows two primer pairs: one for the host (E-a and F-b) and one for microbes (E-c and F-d). Each primer pair adds the same universal overhangs E and F. The PCR reaction (right) shows a single primer pair (P7-E and P5-i-F) that can amplify all tagged products. b, Representative 2% agarose gel of hamPCR products from the synthetic titration panel, showing a V4 16S rDNA amplicon at ~420 bp and an *A. thaliana* GI amplicon at 502 bp. The barplot underneath shows the predicted number of original *GI* and 16S rDNA template copies. Numbers boxed below the barplot indicate the percent bacterial genomic DNA of total DNA. c, Relative abundance of the host and microbial ASVs in the synthetic titration panel, as determined by amplicon counting. Pure *E. coli*, pure *A. thaliana* without PNAs, and blanks were excluded. d, Data in (c) converted to microbial load by dividing by host abundance, with a fourth-root transformed y-axis to better visualize lower abundances.

To assess the technical reproducibility of the protocol, we made a titration panel of artificial samples combining varying amounts of pure *A. thaliana* plant DNA with pure bacterial DNA that reflects a simple synthetic community (Methods). These represented a realistic range of bacterial concentrations as previously observed from shotgun metagenomics of wild leaves, ranging from about 0.25% to 24% bacterial DNA^12^. We applied hamPCR to the panel, pairing one of three commonly-used 16S rDNA amplicons for the V4, V3V4, and V5V6V7 variable regions with either a 502 bp or 466 bp *GI* amplicon (Methods, Supplementary Table 1), such that the host and microbial amplicons differed by approximately 80 bp in length and were resolvable by gel electrophoresis. In all pairings, the *GI* band intensity increased as the 16S rDNA band intensity decreased (Figure 1b, Supplementary Figure 3).

Focusing on the V4 16S rDNA primer set, 515F - 799R, paired with the 502 bp GI amplicon, we amplified the entire titration panel in four independently-mixed technical replicates. In addition to use of the chloroplast-avoiding 799R primer^38^, plant organelle-blocking PNAs^33^ further prevented unwanted 16S rDNA signal from organelles in the pure plant sample (Figure 1b). We pooled the replicates and sequenced them as part of a paired-end HiSeq 3000 lane. Because the 150 bp forward and reverse reads were not long enough to assemble into full amplicons, we analyzed only the forward reads (Methods), processing the sequences into Amplicon Sequence Variants (ASVs) and making a count table of individual ASVs using *Usearch^39^*.

After identifying the ASVs corresponding to host *GI* and the bacteria in the synthetic community, we plotted the relative abundance of *A. thaliana GI,* the three *Sphingomonas* ASVs, and the single *E. coli* ASV across the samples of the titration panel (Figure 1c). There was high consistency between the four replicates, more than what was visually apparent in the gel (Figure 1c, Supplementary Figure 3). We next divided ASV abundances in each sample by the abundance of the host ASV in that sample to give the quantity of microbes per unit of host, a measure of the microbial load. Plotting the data with a fourth-root transformed Y axis, used for better visualization of low bacterial loads, revealed consistent and accurate quantification of absolute microbial abundance from 0 up to about 16% total bacterial DNA (Figure 1d). Through this range, the actual sequence counts for total bacteria matched theoretical expectations based on the volumes pipetted to make the titration (solid black line, Figure 1d). At 16% bacterial DNA, bacteria contributed more than 96% of sequences, and the microbe-to-host template ratio was near 25. At higher microbial loads the trend was still apparent, and the decrease in precision was likely exacerbated by the effects of small numbers; when the host ASV abundance is used as a denominator and the abundance approaches 0, load approaches infinity and all sources of error have a greater influence on the quotient. Through simulations, we decided conservatively that loads calculated with a host ASV abundance below 3% should only be classified as “highly colonized”, and should not be used for quantitative measurements (Supplementary Figure 4). In our case, only a minority of highly infected plants^12^ reached bacterial abundances above this highly-quantitative range.

### hamPCR does not distort the detected composition of the microbial community

We amplified the same wild *A. thaliana* phyllosphere template DNA, either with four technical replicates using V4 16S rDNA primers alone, or alternatively with four technical replicates using hamPCR. After sequencing and deriving ASVs, we first compared ASV abundances within identically-prepared replicates of the pure 16S rDNA protocol to demonstrate best-case technical reproducibility of this established technique. As expected, this resulted in a nearly perfect correlation, with a coefficient of determination R^2^ of 0.99 and abundance distributions as indistinguishable by a Kolmogorov–Smirnov test (Figure 2b). Next, we removed from the hamPCR data the ASV corresponding to *A. thaliana GI*, and rescaled the remaining microbial ASVs to 100% to give relative abundance data. We then compared microbial ASVs from the four pure 16S rDNA replicates to those from the four rescaled hamPCR replicates. In this comparison as well, R^2^ was 0.99 and the distributions were essentially identical (Figure 2c). Thus, the inclusion of a host amplicon in the reaction did not introduce taxonomic biases.

**Figure 2.**
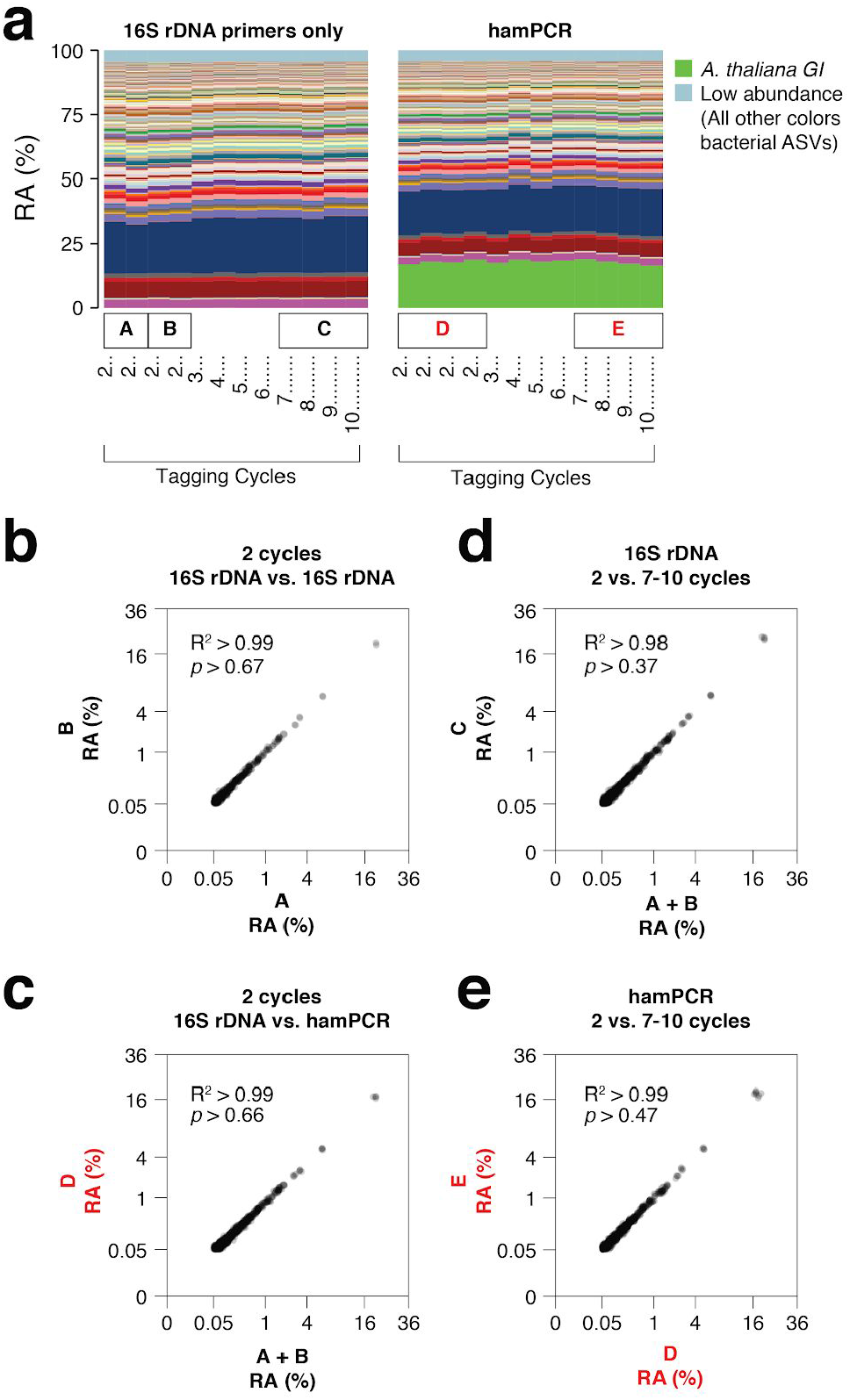
hamPCR is robust and does not distort a complex microbial community. **a**, DNA extracted from wild *A. thaliana* phyllospheres was used as a template for both V4 16S rDNA PCR (left, 515F and 799R) and hamPCR (right, V4 16S rDNA and *GI* 502 bp primers). Four replicates were produced with 2 cycles of the tagging reaction and 30 cycles of PCR, and additional replicates with 3 to 10 tagging cycles paired with 29 to 22 PCR cycles (for a constant total of 32 cycles). The stacked columns show the relative abundances of major ASVs. Boxed upper case letters demarcate groups of samples compared below. **b**, Correlation of fourth-root transformed ASV abundances for the 16S rDNA samples above panel (a) box [A] to the 16S rDNA samples above box [B]. Only ASVs with a minimum relative abundance of 0.05% were compared. R^2^, coefficient of determination. *p*-value from Kolmogorov-Smirnov test. **c**, Same as (b), but for the four 16S rDNA samples above box [A] and [B] compared to the four hamPCR samples above box [D]. For hamPCR, the *A. thaliana GI* ASV was removed and the bacterial ASVs were rescaled to 100% prior to the comparison. **d**, Same as (b) and (c), but for the four 16S rDNA samples above box [A] and [B] compared to the 16S rDNA samples above box [C]. **e**, Same as (b), (c), and (d), **b**ut for the four hamPCR samples above box [D] compared to the four hamPCR samples above box [E].

### Sensitivity to number of tagging cycles and template concentration

Two tagging cycles minimize amplification biases that might otherwise have compounding effects due to differential primer efficiencies for the host and microbial templates. However, for templates at low concentrations, inefficiencies due to SPRI cleanup could represent a bottleneck in amplification. Additionally, some techniques that prevent off-target organelle amplification^40,41^ may benefit from additional tagging cycles. To investigate the sensitivity of the results to additional tagging cycles, we applied hamPCR for 2 through 10 tagging cycles, both on the wild *A. thaliana* phyllosphere DNA described above and on a synthetic plasmid-borne template that contains bacterial rDNA and a partial *A. thaliana GI* gene template in a 1:1 ratio. Surprisingly, for the primers used here, there was no apparent influence of additional tagging cycles, as 7-10 tagging cycles yielded the same distribution of host and 16S rDNA ASV abundances as 2 cycles (Kolmogorov-Smirnov test, *p* > 0.47). This was true for hamPCR and for 16S rDNA primers alone (Figure 2d, 2e). This ideal result may not be the case for all primer pairs and should be tested experimentally, but it is consistent with data that either 5 or 7 tagging cycles gave comparable results for quantifying the human immune receptor repertoire^29^, and with the fact that properly-designed multiplex reactions can be used in qPCR carried out with many cycles^42^. We noticed that application of hamPCR to the 1:1 synthetic template yielded an average of 56.5% host *GI* and 43.5% bacteria, invariant with tagging cycle number (Supplementary Figures 5 and 6). This slight and consistent bias in favor of *GI* may be a result of slight differences in tagging primer efficiency or primer concentration, and should be fine-tunable by altering primer concentration ^29^ (Supplementary Figure 7).

As a further exploration of the robustness of the protocol, we applied hamPCR to a range of total *A. thaliana* leaf template concentrations of between 5 and 500 ng total DNA per reaction, covering a typical template range of 5 to 100 ng. Through the typical range, there was no difference in microbe or host ASV abundances. At 200 ng or above, the host amplicon seemed to be slightly favored, possibly because the 16S rDNA primers started to become limiting at these concentrations (Supplementary Figure 8).

### Pre-sequencing adjustment of host-to-microbe ratio

We realized that the size difference between host and microbe bands in hamPCR affords not only independent visualization of both amplicons on a single gel, but also allows convenient and easy adjustment of the host and microbial signals in the pooled library prior to sequencing, in order to improve cost effectiveness. We developed a strategy by which the final hamPCR amplicons are pooled and one aliquot of the pool is rebarcoded to form a reference sample that preserves the original host- to- microbe ratio. The remainder of the pool is run on a gel and the host and microbial bands are separately purified, quantified, and remixed (in our case, to reduce host and gain more microbial resolution). The rebarcoded reference sample, which was not remixed and thereby preserves the original ratio, can be sequenced separately or spiked into the remixed library prior to sequencing. Following sequencing, the reference sample provides the key to the correct host and microbe proportions, allowing simple scaling of the entire library back to original levels (Figure 3a, Supplementary Figure 9).

**Figure 3.**
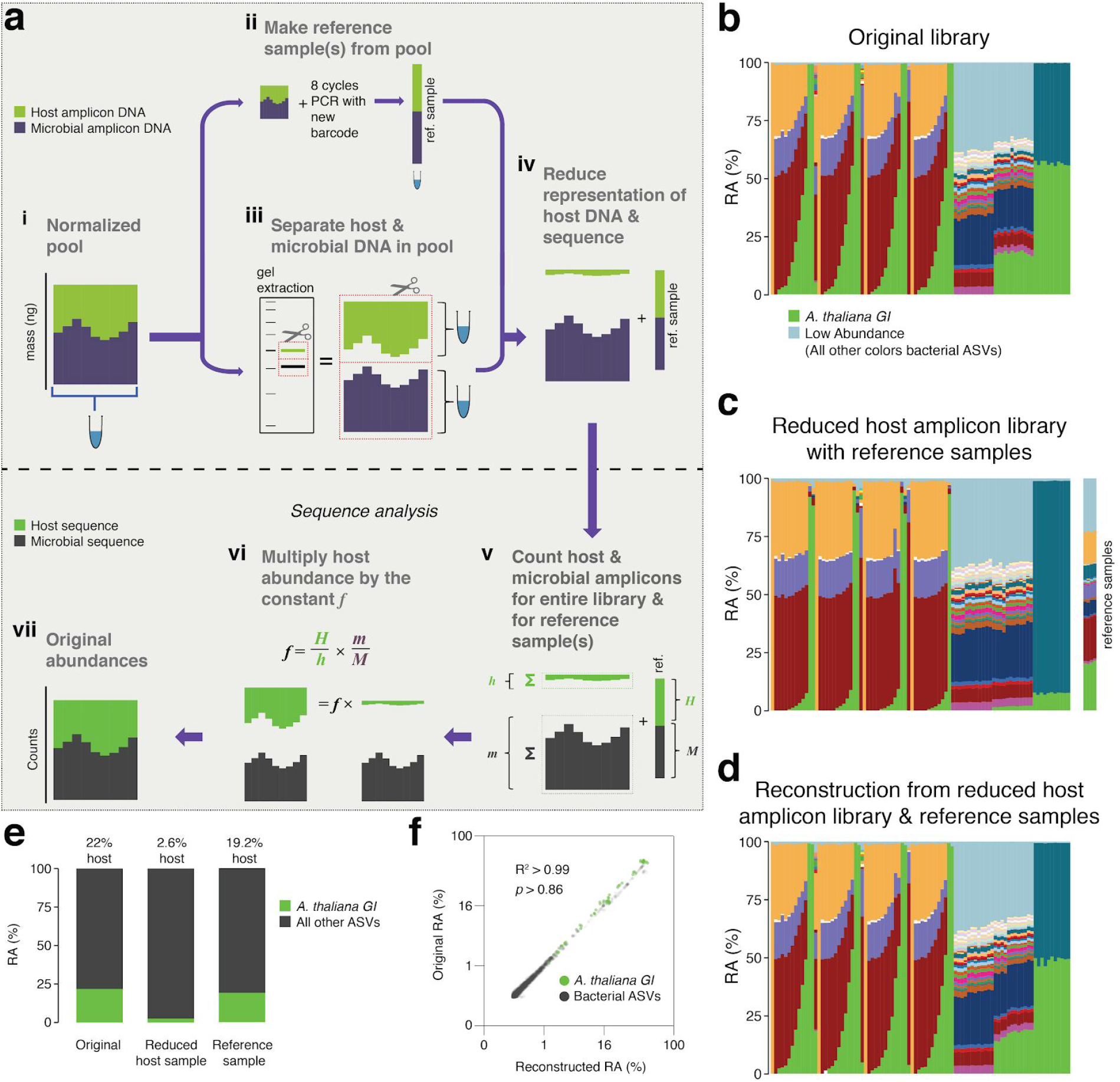
After remixing hamPCR amplicons for efficient sequencing, original abundances can be reconstructed. **a**, Scheme of remixing process. **i**: Products of individual PCRs are pooled at equimolar ratios into a single tube. **ii**: An aliquot of DNA from the pool in (**i**) is re-amplified with 8 cycles of PCR to replace all barcodes in the pool with a new barcode, creating a reference sample. **iii**: An aliquot of the pool from (**i**) is physically separated into host and microbial fractions via agarose gel electrophoresis. **iv**: The host and microbial fractions and the reference sample are pooled in the ratio desired for sequencing. **v**: All sequences are quality filtered, demultiplexed, and taxonomically classified using the same parameters. **vi**: Host and microbial amplicon counts are summed from the samples comprising the pooled library (*h* and *m* respectively), and from the reference sample (*H and M*). **vi**: *H, h, M, and m* are used to calculate the scaling constant *f* for the dataset. All host sequence counts are multiplied by *f* to reconstruct the original microbe-to-host ratios. **vii**: Reconstructed original abundances. **b**, Relative abundance (RA) of actual sequence counts from our original HiSeq 3000 run. **c**, Relative abundance of actual sequence counts from our adjusted library showing reduced host and 4 reference samples. **d**, The data from (c) after reconstructing original host abundance using the reference samples. **e**, The total fraction of host vs. other ASVs in the original library, reduced host library, and reconstruction. **f**, Relative abundances in the original and reconstructed library for all ASVs with a 0.05% minimum abundance, shown on 4^th^-root transformed axes. R^2^, coefficient of determination. *p*-value from Kolmogorov–Smirnov test.

We made 4 replicate reference samples for our HiSeq 3000 run, which included much of the data from Figure 1 and Figure 2, and then separately purified the host and microbial fractions of the library (Supplementary Figures 9 and 10). Based on estimated amplicon molarities of the host and microbial fractions, we remixed them targeting 5% host DNA, added the reference samples, and sequenced the final mix as part of a new HiSeq 3000 lane. A stacked-column plot of relative abundances for all samples on the original run clearly showed the host *A. thaliana GI* ASV highly abundant in some samples, on average responsible for about 22% of total sequences in the run (Figure 3b, 3e). The remixed reduced host library had nearly 10-fold less host *GI* ASV, 2.6%, slightly lower than our target of 5% (Figure 3c, 3e). The reference samples averaged 19.2% of host *GI* ASV, very close to the 22% host fraction in the original library. After using the reference samples to reconstruct the original host abundance in the remixed dataset, we recreated the shape of the stacked column plot from the original library (compare Figure 3d to 3b). When the fourth-root abundances for ASVs above a 0.05% threshold were compared between the original and reconstructed libraries, the R^2^ coefficient of determination was 0.99, with no significant difference between the distributions (Kolmogorov-Smirnov test, *p* > 0.86).

### hamPCR with different 16S rDNA regions compared to shotgun metagenomics

We next applied hamPCR to leaf DNA from eight wild *A. thaliana* plants that we had previously analyzed by WMS, and from which we therefore had an accurate estimate of the microbial load as the number of microbial reads divided by the number of plant chromosomal reads^12^. We applied hamPCR with primer combinations targeting the host *GI* gene and either the V3V4, V4, or V5V6V7 variable regions of the 16S rDNA. We produced three independent replicates for each primer set, which we averaged for final analysis (Figure 4, Supplementary Figure 11). Across WMS and the three hamPCR amplicon combinations, the relative abundance of bacterial families was consistent (Figure 4a-d: i), with slight deviations likely due to the different taxonomic classification pipeline used for the metagenome reads^12^, as well as known biases resulting from amplification or classification of different 16S rDNA variable regions^43^. After converting both WMS and hamPCR bacterial reads to load by dividing by the plant reads in each sample, we recovered a similar pattern despite the quantification method, with decreasingly lower total loads progressing from plant S1 to plant S8, and individual bacterial family loads showing similar patterns (Figure 4a-d: ii, iii). Relative differences in load estimates when comparing the different hamPCR amplicons are likely in part due to different affinities of the 16S rDNA primer pairs for their targets in different bacterial species, and rDNA copy number variation among the microbial families^44^. To quantify the consistency of hamPCR load estimates with WGS load estimates, we plotted the loads against each other and found strong positive correlations, with the highest correlation with the hamPCR using V4 rDNA (Figure 4e-g).

**Figure 4.**
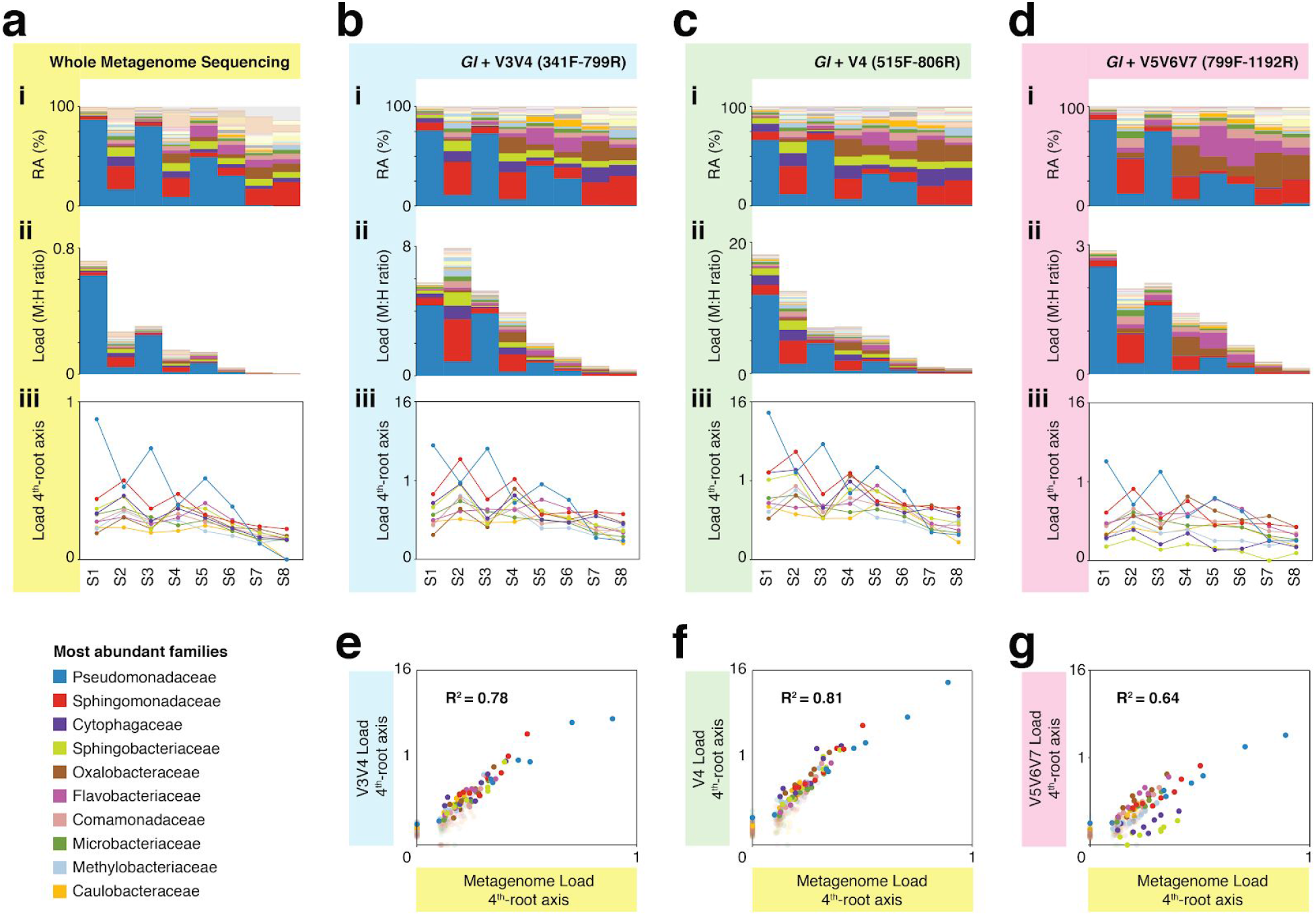
hamPCR with three common 16S rDNA amplicons gives consistent results that agree with shotgun metagenomics. **a**, **i**: Stacked-column plot showing the relative abundance (RA) of bacterial families in eight wild *A. thaliana* leaf samples, as determined by shotgun sequencing. The families corresponding to the first 10 colors from bottom to top are shown in reverse order on the bottom left. **ii**: Stacked-column plot showing the bacterial load of the same bacterial families (M:H ratio = microbe-to-host ratio). **iii**: The M:H bacterial load ratios for the 10 major bacterial families shown on a 4^th^-root transformed y-axis. Lines across the independent samples are provided as a help to visualize patterns. **b**, Similar to (a), but with abundances resulting from hamPCR targeting a 502 bp *A. thaliana GI* amplicon and a ~590 bp V3V4 16S rDNA amplicon. **c**, Similar to (b), but with the 16S rDNA primers targeting a ~420 bp V4 16S rDNA amplicon. **d**, Similar to (b), but with a 466 bp *A. thaliana GI* amplicon and a ~540 bp V5V6V7 16S rDNA amplicon. **e**, 4^th^root transformed abundance of each bacterial family determined by hamPCR of V3V4 16S rDNA plotted against the 4^th^-root transformed bacterial load from shotgun metagenomics. R^2^ = Coefficient of determination. **f**, Same as (e), but for hamPCR of V4 16S rDNA. **g**, Same as (e), but for V5V6V7 16S rDNA.

It is important to note that while relative load ratios between samples were consistent across hamPCR primer sets, the total microbe-to-host ratio varied substantially, with the maximum V5V6V7 16S rDNA total load at less than 3 times host, and the maximum V4 16S rDNA total load near 16 times host. This is likely due to variation in *GI* and 16S rDNA primer efficiencies. To make a statement about the ratio of plant cells to bacterial cells using hamPCR, it would be important to include standard samples with known bacterial load ratios, and to normalize each bacterial taxon by its average rDNA copy number.

### Three-amplicon hamPCR for simultaneous determination of oomycete and bacterial load

More than two amplicons can be quantitatively tagged and amplified, although it requires more initial troubleshooting to find compatible primers^29,30,42,45^. We set up hamPCR to co-quantify bacteria and eukaryotic oomycetes on plants. These are both diverse microbial groups that include important *A. thaliana* pathogens and that cannot be captured by the same rDNA primer set. We first tested universal ITSo primers targeting oomycete rDNA (Methods) in combination with *A. thaliana GI* primers and 16S rDNA primers targeting the bacterial V4, V3V4, or V5V6V7 regions, using as template our synthetic plasmid that includes templates for the three primer sets in equal proportion (Methods). A combination of all three amplicons seemed to work efficiently for the V4 region (Supplementary Figure 12), and with this encouraging result, we set up a simple infection experiment. As pathogens, we prepared local strain 466-1 of the obligate biotrophic oomycete *Hyaloperonospora arabidopsidis* (*Hpa*)^46^ and the well-described bacterial pathogen *Pseudomonas syringae* pv. *tomato* (*Pto*) DC3000 ^47^. We used two *A. thaliana* genotypes: the reference accession Col-0, which is resistant to *Hpa* 466-1 but susceptible to *Pto* DC3000, and an *enhanced disease susceptibility 1* (*eds1-1*) mutant, which has a well-studied defect in a lipase-like protein necessary for many disease resistance responses and which is susceptible to both pathogens^48^.

We infected seedlings with either *Hpa* 466-1 alone, a mix of *Hpa* 466-1 and *Pto* DC3000, or a buffer control, and maintained them for 7 days under cool, humid conditions ideal for *Hpa* growth (Methods). The *eds1-1* plants inoculated with *Hpa* 466-1 became heavily infected and sporangiophores were too numerous to count. No visible bacterial disease symptoms were present on any of the plants, likely because the cool temperature decelerated bacterial growth and symptom appearance^49^. We ground pools of 4-5 seedlings in a buffer and used a small aliquot to count *Pto* DC3000 CFUs, and the remainder of the lysate for DNA isolation and hamPCR. Despite the lack of bacterial symptoms, we recovered *Pto* DC3000 CFUs from the inoculated plants.

We applied hamPCR to these samples using the ITSo/16S/*GI* primer set, but due to excessive ITSo product, we repeated library construction replacing the ITSo primers with primers for a single copy *Hpa* actin gene (Supplementary Figure 12)^23^. Intensity of the actin product correlated with visual *Hpa* symptoms (Supplementary Figure 12). Sequencing the libraries confirmed *Hpa* and *Pto* ASVs in the inoculated samples, as expected. A standard bacterial relative abundance plot, as would be obtained from pure 16S rDNA data, confirmed the presence of *Pto* DC3000 in the bacteria-infected samples, and in addition revealed that *Hpa*-infected samples had a different bacterial community than uninfected samples (Figure 5a). Importantly, it failed to detect obvious differences between microbial communities on Col-0 and *eds1-1* plants. However, after including the actin ASV from *Hpa* and converting all abundances to microbial load, a striking difference became apparent between Col-0 and *eds1-1*, with *eds1-1* supporting higher bacterial and *Hpa* abundances. This is expected from existing knowledge^48^, and supported by *Pto* DC3000 CFU counts from the same plants (Figure 5b, Figure 5c). The microbial load plot also revealed that *Hpa*-challenged plants supported more bacteria than buffer-treated plants, indicating either that successful bacterial colonizers were unintentionally co-inoculated with *Hpa*, or that *Hpa* caused changes in the native flora (Figure 5b).

**Figure 5.**
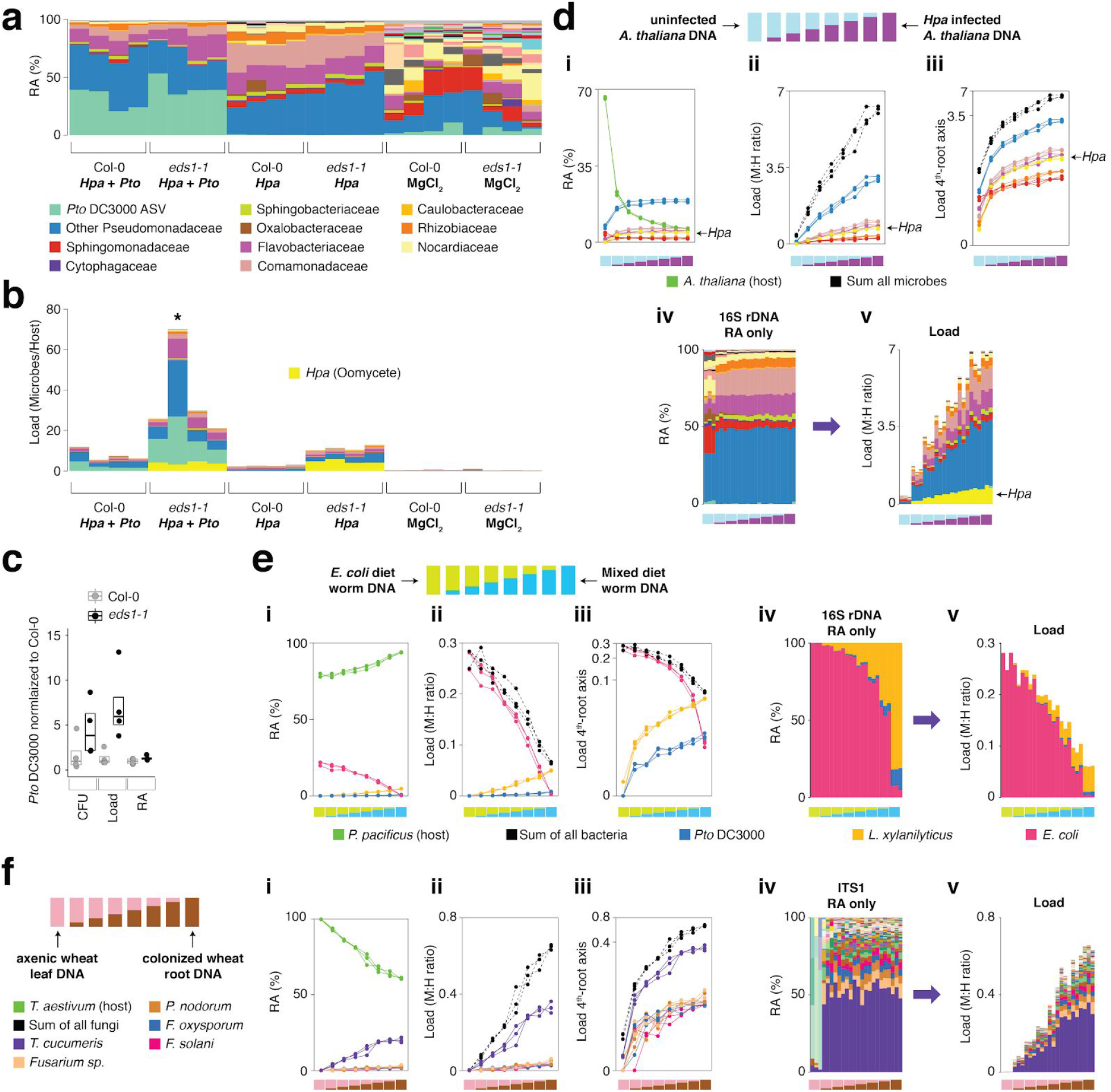
hamPCR can be generalized to more than two amplicons, non-plant hosts, and large host genomes. **a**, Relative abundance (RA) of only 16S rDNA amplicons for plants co-infected with *Hpa* and *Pto* DC3000 and their controls. The ASV corresponding to *Pto* DC3000 (light green) is shown at the bottom and separately from other Pseudomonadaceae; all other bacteria are classified to the family level, including remaining Pseudomonadaceae. **b**, The same data as shown in (a), but converted to microbial load and with the ASV for *Hpa* included. The asterisk indicates what is most likely an unreliable load calculation, because host ASV abundance was below 3% of the total. Same color key as in (a), with an additional color (yellow) for *Hpa* added. *Hpa* amplicon abundance was scaled by a factor of 4 in this panel for better visualization. **c**, *Pto* DC3000 bacteria were quantified in parallel on the Col-0 and *eds1-1* samples infected with *Pto* DC3000 using CFU counts, the microbial load data in (b), or the relative abundance data in (a). The median is shown as a horizontal line and box boundaries show the lower and upper quartiles. **d**, An uninfected plant sample was titrated into an *Hpa*-infected sample to make a panel of eight samples. **i**: the relative abundance of hamPCR amplicons with median abundance above 0.15%. **ii**: after using host ASV to convert amplicons to load. The cumulative load is shown in black. **iii**: the load on a 4^th^-root transformed y-axis, showing less-abundant families. **iv**: stacked column visualization of all ASVs for the panel as it would be seen with pure 16S rDNA data. **v**: stacked-column plot of the panel corrected for microbial load. Same color key as in (b), but with colors for *A. thaliana* and sum of microbes added. **e**, Similar to (d), but with the nematode worm *P. pacificus* as host, and V5V6V7 16S rDNA primers. Instead of bacterial families, specific ASV abundances are shown. **f**, Similar to (e), but with hexaploid wheat *T. aestivum* as host, and fungal ASV abundances from ITS1 amplicons.

To confirm that the sequence abundances for all three amplicons accurately reflected the concentration of their original templates, we prepared a stepwise titration panel with real samples, mixing increasing amounts of DNA from an uninfected *eds1-1* plant (low load) into decreasing amounts of DNA from an *Hpa*-infected *eds1-1* plant (high load). Sequencing triplicate hamPCR libraries revealed a stepwise increase in ASV levels for all amplicons, consistent with the expectation based on pipetting (Figure 5d). These data, combined with the infection experiment, show that hamPCR is quantitative for at least two independent microbial amplicons in real-world samples.

### Utility in diverse hosts and crops with large genomes

To demonstrate the utility of hamPCR outside of plants, we prepared samples of the nematode worm *Pristionchus pacificus*, fed on a diet of either pure *E. coli* OP50, or alternatively a mix of *E. coli* OP50 with *Pto* DC3000 and *Lysinibacillus xylanilyticus*. The worms were washed extensively with PBS buffer to remove epidermally-attached bacteria, enriching the worms for gut-associated bacteria, and we prepared DNA from each sample. In the same manner as described in the previous section, we titrated the two DNA samples into each other to create a panel of samples representing a continuous range of colonization at biologically-relevant levels. Over three replicates, hamPCR accurately captured the changing bacterial loads of the gut microbes (Figure 5e, Supplementary Figure 13). We similarly validated the technique for fungal and bacterial microbes of *Triticum aestivum* (bread wheat) the most widely grown crop in the world and one of the most difficult to study due to a 16 Gb haploid genome size^50^. To simulate different levels of infection, we titrated DNA from axenically-grown wheat leaves into DNA from wheat roots that had been cultivated in non-sterile soil and applied hamPCR, using as a host gene RNA polymerase A1 (*PolA1*), which is present as a single copy in each of the A, B, and D subgenomes^51^. We recovered expected abundance patterns in the panel both for ITS1 rDNA primers (Figure 5f, Supplementary Figures 14 and 15) and for V4 16S rDNA primers (Supplementary Figures 14 and 15). We noticed the original ratio of ITS1 to *PolA1* sequences recovered was low; because ITS primers produce amplicons that are highly variable in length, some of which may co-migrate with the host amplicon on a gel, the cut-and-mix approach described in Figure 3 could not be used to improve ITS1 representation. However, increasing the ratio of the ITS1:*PolA1* tagging primers from 1:1 to 2:1 (Methods) successfully enriched the ITS1 amplicon without sacrificing relative load determination between samples (Supplementary Figures 14 and 15).

To demonstrate the ability of hamPCR to yield new biological insights into complex study systems, we conducted two experiments with crop plants. First, we set up a growth curve in bell pepper (*Capsicum annuum*), which has a 3.5 Gb genome^52^, approximately 25× larger than *A. thaliana*, and the pepper pathogenic bacterium *Xanthomonas euvesicatoria* (*Xe*) strain 85-10^53^. As proof-of-concept preparation for the growth curve, to confirm that hamPCR could accurately capture absolute changes in pathogen abundance in pepper leaves, we constructed an infiltration panel in which *Xe* 85-10 was diluted to final concentrations of 10^4^, 10^5^, 10^6^, 10^7^ and 10^8^ CFU / mL and infiltrated into four replicate leaves per concentration. Immediately afterwards, without further bacterial growth, we harvested leaf discs within inoculated areas using a cork borer. We ground the discs and used some of the lysate for *Xe* 85-10 CFU counting, and the remainder for DNA extraction and hamPCR targeting the V4 16S rDNA and the pepper *GI* gene (*CaGI*), and qPCR, targeting the *xopQ* gene for a *Xe* type III effector ^54^, and the *C. annuum UBI-3* gene for a ubiquitin-conjugating protein^55^.

Sequencing the hamPCR libraries revealed that as the *Xe* 85-10 infiltration concentration increased, so did the resulting load of the ASV corresponding to *Xe* 85-10 (Supplementary Figure 16 and 17a). The other major bacterial classes detected in the infiltration panel, comprising commensal bacteria already present in the leaves, had similar, low abundances, regardless of the amount of infiltrated *Xe* 85-10 (Supplementary Figure 17b). When aligned with CFU counts recovered from the same lysates (Methods), the hamPCR *Xe* 85-10 ASV loads showed nearly the same exponential differences between samples, although at lower infiltration concentrations, qPCR and hamPCR gave a slightly higher estimate than CFU counts (Supplementary Figure 17c). The presence of a low level of native, antibiotic-sensitive *Xe* on the leaves could potentially explain this discrepancy, because this could be detected by DNA-based methods but not culturing.

For the pepper growth-curve, we infiltrated six *C. annuum* leaves of six different plants with *Xe* 85-10 at a concentration of 10^4^ CFU / mL, and took samples from each plant at 0, 2, 4, 7, 9 and 11 dpi for CFU counting, qPCR, and hamPCR. We observed a rapid increase in *Xe* 85-10 ASV abundance as a result of rapid bacterial growth, leveling off at 7 dpi (Figure 6a). By 7 dpi, bacterial growth had reduced the host *GI* amplicon abundance to below 3%, making the load corrections unreliable from day 7 on (gray box, Figure 6a). Aligned *Xe* 85-10 ASV loads compared very closely to CFU counts and to aligned qPCR abundances up to 7 dpi (Figure 6c). Notably, the other major bacterial classes, Actinobacteria and the Alpha-, Beta-, and Gammaproteobacteria, also increased in microbial load through time, a trend significant even comparing 2 dpi to 0 dpi (Figure 6b, Mann-Whitney U-test, *p* < 0.001). This increase in load for the other classes was not a PCR artifact due to high *Xe* 85-10 titers, because in the infiltration panel, measurements for these classes had not changed even at higher pathogen concentrations (Supplementary Figure 17b). This subtle but biologically significant effect of infection on growth of commensal bacteria would be completely invisible in a pure 16S rDNA amplicon analysis, which would only show *Xe* 85-10 overtaking the community.

**Figure 6.**
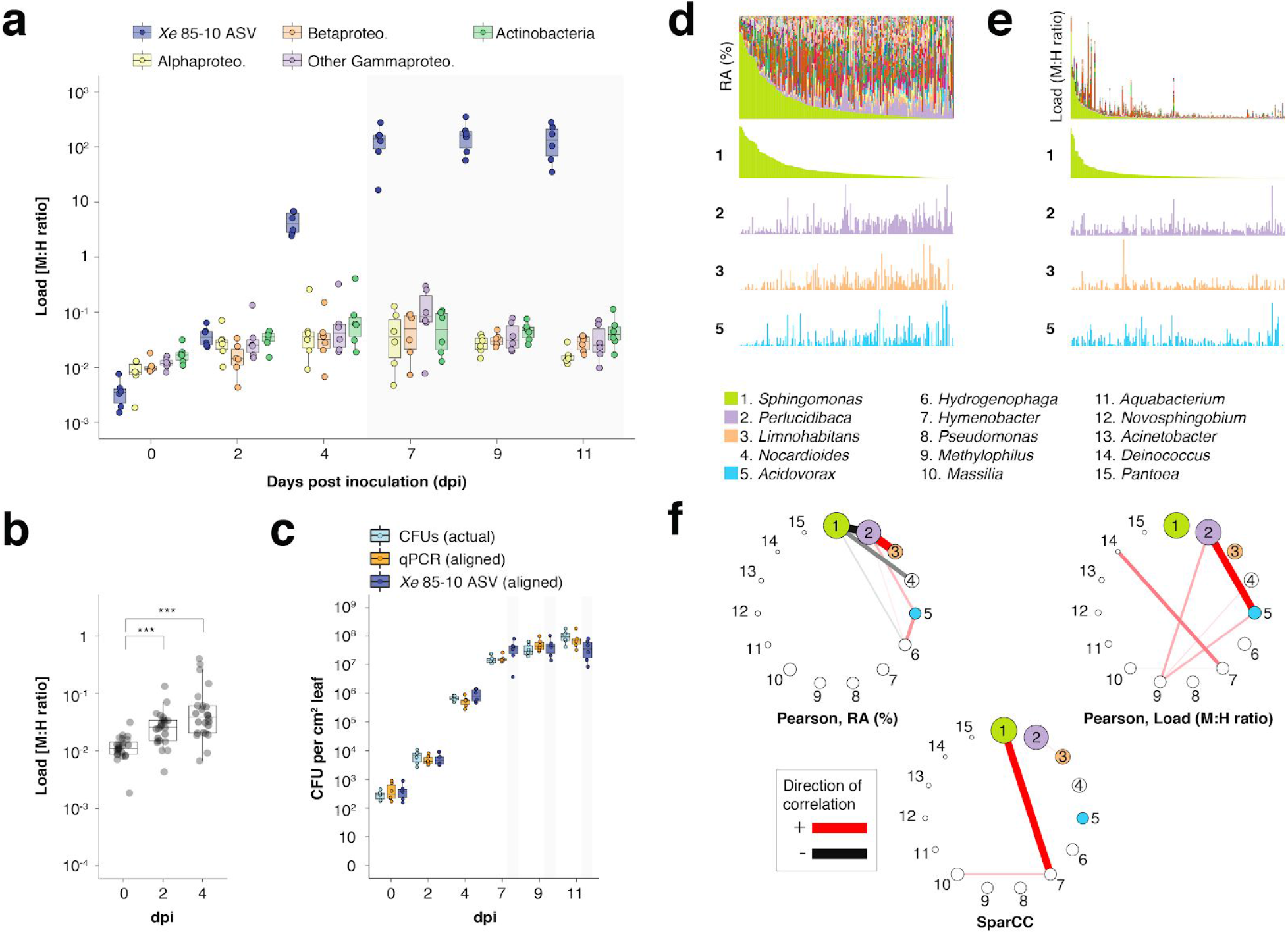
hamPCR can provide new insights into microbial interactions in crop plants. **a-c**, *C. annuum* growth curve experiment. All y-axes are on a base-10 logarithmic scale. In all boxplots, the median is represented by a horizontal line and box boundaries show the lower and upper quartiles. Whiskers extend from the box up to 1.5 times the interquartile range. **a**, *Xe* 85-10 was inoculated into *C. annuum* leaves at 104 CFU/mL. Leaf samples were taken at 0, 2, 4, 7, 9, and 11 days post inoculation (dpi), and hamPCR performed. The corrected load is shown for the particular ASV corresponding to *Xe* 85-10, as well as for the major bacterial classes. **b**, The total load for all bacterial classes shown in (d) at 0, 2, and 4 dpi (*** = *p* < 0.001). **c**, Actual CFU counts for *Xe* 85-10 in the growth curve experiment juxtaposed with aligned qPCR and hamPCR loads. **d-e**, Field-grown *Z. mays* collection. **d**, Relative abundance (RA) of bacterial genera found in *Z. mays* leaf hole punches, ordered by *Sphingomonas* relative abundance. The genera corresponding to the first 15 colors from bottom to top are shown in reverse order in the legend. The relative abundance of four isolated genera is highlighted (colored boxes in legend). **e**, same as **d**, but showing microbial load rather than relative abundance and ordered by *Sphingomonas* load. **f**, Correlation networks of the same 15 genera from the legend for **d** and **e**. Pearson correlation from RA data from **d** (left), pearson correlation of microbial load from **e** (right), and SparCC correlation network (bottom). Circles representing genera are scaled such that their area represents the median genus abundance across all samples. Only correlations of absolute magnitude >= 0.3 are shown.

Finally, we applied hamPCR to DNA from 201 leaf samples from mature, isogenic maize (B73) growing in a field site in Tübingen, Germany. We used the V4 region of 16S rDNA for bacteria and the single copy *LUMINIDEPENDENS* (*LD*) gene as a host marker, and plotted both the relative abundance of bacterial genera (Figure 6d) and the bacterial load of these genera (Figure 6e). In some samples, the genus *Sphingomonas* exceeded 80% of the bacterial community, creating especially strong compositionality effects; other abundant genera *Perlucidibaca, Limnohabitans,* and *Acidovorax* visibly increased in relative abundance as *Sphingomonas* became less abundant (Figure 6d). In contrast, the bacteria load of these same genera appeared mostly unaffected by *Sphingomonas* bacterial load (Figure 6e). As expected, a Pearson correlation network made with relative abundance data revealed that *Sphingomonas* was negatively correlated with many genera, a well-known and problematic artifact of compositionality^56^ (Figure 6f). A Pearson correlation network made with microbial load data was remarkable in that *Sphingomonas,* despite having the highest median abundance of any genus, is among the genera least correlated with others (Figure 6f). We also calculated a correlation network using SparCC^56^, which estimates Pearson correlations on log-transformed components to avoid compositionality artifacts. This network did indeed avoid the spurious negative correlations with *Sphingomonas*, although it still implicated the genus more strongly than the true correlation network built with hamPCR data. Each network has a very different biological interpretation. If *Sphingomonas* abundance does not influence these other genera on healthy leaves, this could mean, for example, that it can colonize more of the available extreme habitat, with its success determined largely by abiotic factors. Future study will be necessary to resolve this. Overrepresentation of *Sphingomonas* is a feature shared by other major studies of the maize leaf microbiome^57,58^; overcoming this compositionality problem is broadly relevant to studies of this microbial habitat.

## Discussion

We developed hamPCR, a simple and robust method to quantitatively co-amplify one or more microbial marker genes along with an unrelated host gene, allowing accurate determination of microbial load and microbial community composition from a single sequencing library (Figure 1, Figure 2). Furthermore, we developed a method to predictably optimize the amount of sequencing effort devoted to microbe vs. host, without losing information about the original microbe to host ratio (Figure 3). This is an important advance in our approach that greatly increases cost-efficiency.

The principle behind hamPCR stands on a body of literature describing related, firmly established techniques, which bodes well for wide-spread adoption of our approach. Using two steps in a PCR protocol is common in amplicon sequencing, including of microbial marker genes^31–33,59,60^. Two-step PCR protocols provide the major advantage that only a one-time investment is needed in a set of universal barcoding primers for a flexible step two. These can be easily adapted to any amplicon(s) by simply swapping in different template-specific primers for step one. For labs already equipped for two-step PCR, implementing hamPCR involves only slight adjustments to cycling conditions and template-specific tagging primers.

Quantitative co-amplification using multiple primer pairs also has proven reliable^29,30,61^, and PCR biases affecting co-amplification of diverse fragments are manageable and well-understood from popular RNA-seq and WMS protocols^62,63^. This rich literature should increase confidence when implementing hamPCR in microbiome research, and it also provides resources for optimization and further development. For example, the use of fewer cycles in exponential PCR could reduce noise and bias, hamPCR tagging primers could be fitted with UMIs for higher precision, and the protocol could be adapted for sequencing platforms with longer read lengths.

We have demonstrated that microbial load measurement is sensitive to the relative concentrations between the host and microbe primers in the tagging step (Supplementary Figure 7), consistent with the effects of primer concentration on amplification efficiency in qPCR^64–66^. This property makes it possible to fine-tune the primer ratios, either to yield the expected ratio of products^29^, or to intentionally increase the representation of a microbial amplicon for more efficient sequencing (Supplementary Figure 16a). The effect of primer concentration has important implications for how a large project should be prepared. We recommend that the tagging primers be carefully pipetted into a multiplexed primer master mix sufficiently large to be used for the entire project, or alternatively the same control samples should be sequenced across sample batches to allow correction of slight batch differences.

A limitation of hamPCR is reduced accuracy at the highest microbial loads (Supplementary Figure 4). Only a minority of our samples reached a level of infection that interfered with accurate quantification, and we expect that this will be the case for most colonized hosts. If not, there are three straightforward adjustments that can increase host signal to acceptable levels. First, altering the host and microbe amplicon ratio in the pooled library prior to sequencing, as demonstrated in Figure 3, could be used to *increase* the overall host representation. Second, a host gene with a higher copy number could be chosen for template tagging throughout the entire project. Finally, adjusting the concentration of the host primers in the tagging reaction could also increase the representation of host (Supplementary Figure 7)^29^.

In summary, we have demonstrated that hamPCR is agnostic to the taxonomic identities of the organisms studied on both the host and microbe side, their genome sizes, or the functions of the regions amplified. We have also shown that hamPCR can monitor three amplicons at the same time for interkingdom microbial quantification, and in principle can multiplex more with careful design. Our focus here has been on tracking hosts and their closely-associated microbes, but the protocol could also be adapted to quantitatively relate different amplicons targeting archaea, bacteria, and fungi in diverse “host-free” environments like soil. Besides whole organisms, hamPCR also enables quantitative monitoring of bacterial populations and sub-genomic elements, such as plasmids or pathogenicity islands that might not be shared by all strains in a population. An exciting application of hamPCR is the study of endophytic microbial colonization and infection in crop plants, many of which have very large genomes that preclude the analysis of any sizable number of samples by shotgun sequencing. In a previous study, we sequenced leaf metagenomes from over 200 *A. thaliana* plants, at not insignificant costs^12^. In wheat, assuming comparable microbial loads, the same investment in sequencing would barely be sufficient for two samples due to the size of the wheat genome of over 16 Gb. Microbial analysis of the >200 samples we processed of field-grown maize likewise would be prohibitively expensive by shotgun sequencing, and supplementing these data with an orthogonal method on this scale requires at least double the sample and double the time.

Other exciting applications are the recognition of cryptic infections^67^, tracking of mixed infections, and measurement of pathogen abundances on hosts showing quantitative disease resistance - this could even be accomplished by spiking hamPCR amplicons into the same sequencing run used to genotype the hosts^68^. In sparsely-colonized samples, hamPCR will help prevent inflating the abundance of ultra-low abundance microbes, such as reagent contaminants. Finally, for projects with many samples, the fact that hamPCR derives microbial composition and load from the same library not only saves costs and uses less of the sample, but also simplifies analysis and project organization.

## Methods

### hamPCR protocol

hamPCR requires two steps: a short ‘tagging’ reaction of 2 cycles, and a longer ‘exponential’ reaction. We used 30 cycles throughout this work, although fewer can and should be used if the signal is clear for better quantitative results. The primers employed in the tagging reaction were used at ⅛ the concentration of the exponential primers, as this still represents an excess in a reaction run for only two cycles, prevents waste, and reduces dimer formation. See Supplementary Information and Supplementary Table 1 for detailed information about the primers.

#### Tagging reaction

We used Taq DNA Polymerase (NEB, Ipswich, MA, USA) for the first tagging step, and set up 25 μL reactions as follows.

| Master Mix | For 25 μL |
| --- | --- |
| 10x Taq buffer | 2.5 μL |
| Taq polymerase | 0.2 μL |
| dNTPs (10 mM) | 0.5 μL |
| * Tagging primer mix | 1.25 μL |
| ** PNAs (mix of mPNA and pPNA each at 50 μM) | 0.375 μL |
| *** DNA (2-10 ng/μL) | 5 μL |
| Water | (to 25 μL) |

\* tagging primer mix is an equimolar mix of tagging primers with each at a partial concentration of 1.25 μM
\*\* PNA was used to block chloroplast and mitochondrial amplification in reactions involving the V4 or V3V4 region of 16S rDNA. For other reactions, it is not helpful and is omitted.
\*\*\* not part of master mix

Each well received 20 μL of master mix and 5 μL of DNA (around 50 ng). Completed reactions were thoroughly mixed on a plate vortex and placed into a preheated thermocycler. We used the following standard cycling conditions:

1. **94° C** for 2 min. ***Denature***
2. **78° C** for 10 sec. ***PNA annealing***
3. **58° C** for 15 sec. ***Primer annealing***
4. **55° C** for 15 sec. ***Primer annealing***
5. **72° C** for 1 min. ***Extension***
6. **GO TO STEP 1** for **1** additional cycle
7. **16° C** forever ***Hold***

The tagging reaction was cleaned with Solid Phase Reversible Immobilization (SPRI) beads^36^. All ITS amplicons were cleaned with a 1.1:1 ratio of SPRI beads to DNA, or 27.5 μL beads mixed in 25 μL of tagged template. After securing beads and DNA to a magnet and removing the supernatant containing primers and small fragments, beads were washed twice with 80% ethanol, air dried briefly, and eluted in 17 μL of water. For primer sequences, see Supplementary Information and Supplementary Table 1.

#### Exponential reaction

15 μL of the tagged DNA from step one was used as template for the exponential reaction. To reduce errors during the exponential phase, we used the proof-reading enzyme Q5 from NEB, with its included buffer. We prepared reactions in 25 μL for technical tests with replicated samples. For samples prepared without sequenced replicates, we prepared most in triplicate reactions in which a 40 μL mix was split into 3 parallel reactions of ~13 μL prior to PCR to reduce bias, although this is likely unnecessary ^69^.

| Master Mix | For 25 μL | For 40 μL |
| --- | --- | --- |
| 5x Q5 buffer | 5 μL | 8 μL |
| Q5 polymerase | 0.25 μL | 0.4 μL |
| dNTPs (10 mM) | 0.5 μL | 0.8 μL |
| 100 μM F universal PCR primer | 0.0625 μL | 0.1 μL |
| * PNAs (mix of mPNA and pPNA each at 50 μM) | 0.375 μL | 0.6 μL |
| ** 5 μM reverse barcoded primer | 1.25 μL | 2 μL |
| ** DNA (from previous reaction) | 15 μL | 15 μL |
| Water | (to 25 μL) | (to 40 μL) |

\* PNA was used to block chloroplast and mitochondrial amplification in reactions involving the V4 or V3V4 region of 16S rDNA. For other reactions, it is not helpful and is omitted.
\*\* not part of master mix

We first distributed 8.75 μL (or 23 μL for 40 μL mixes) of master mix to each well. We then added 15 μL of the DNA from the tagging reaction and 1.25 μL (or 2 μL for 40 μL mixes) of 5 μM barcoded reverse primer. For the 40 μL mixes, 13 μL was pipetted into two new PCR wells. The PCR reactions were placed into a hot thermocycler and cycled with the following standard conditions:

1. **94° C** for 2 min. ***Denature***
2. **94° C** for 20 sec. ***Denature***
3. **78° C** for 5 sec. ***PNA annealing***
4. **60° C** for 30 sec. ***Primer annealing***
5. **72° C** for 45 sec. ***Extension***
6. **GO TO STEP 2** for **29** additional cycles
7. **16° C** forever ***Hold***

Following PCR, sets of three 13 μL reactions were recombined to 40 μL. For primer sequences, see Supplementary Information and Supplementary Table 1.

### Library quality control and pooling

For visualization, 5 μL of PCR product was mixed with 3 μL of 6x loading dye and all 8 μL loaded on a 2% agarose gel and stained with ethidium bromide. The remaining PCR products were cleaned with a SPRI-to-DNA ratio of 1.1:1.0 (v/v). The DNA concentrations in the cleaned products were measured with PicoGreen (Invitrogen, Carlsbad, CA, USA) and samples were pooled at equimolar total DNA ratios. We note that because host and microbial fractions are independently visible on the gel, it would also be possible to measure the quantity of microbial products with image analysis software such as ImageJ^70^ and pool at equimolar microbial ratios.

The pooled library was diluted to ~1 ng/uL and run on a Bioanalyzer High Sensitivity DNA chip (Agilent, Santa Clara, CA, USA) to check library purity and to estimate the expected ratio of host to microbial amplicons in the sample.

### Pre-sequencing adjustment of host : microbe Ratio

To adjust the host-to-microbe ratio in the “synthetic template panel” and “cycle number test” prior to sequencing on a HiSeq3000 instrument (Illumina, San Diego, CA, USA), four reference samples were first made by rebarcoding the original pooled library (Supplementary Figures 9 and 10). To accomplish this, ~5 ng of of the pooled library was used in a 30 μL PCR reaction as follows:

| Master Mix | For 30 μL |
| --- | --- |
| 5x Q5 buffer | 6 μL |
| Q5 polymerase | 0.3 μL |
| dNTPs (10 mM) | 0.6 μL |
| 100 μM F universal PCR primer | 0.075 μL |
| ** 5 μM reverse barcoded primer | 1.5 μL |
| ** 5 ng original pooled library | 5 μL |
| Water | 16.45 μL |

\*\* not part of master mix

After distributing 23.5 μL of master mix to each well, 5 μL of the diluted original library was added to each well (5 ng total), along with 1.5 μL of 5 μM barcoded reverse primer. Just prior to placing the reactions in the thermocycler, a 5 μL pre-PCR aliquot was removed from each one and kept on ice to preserve the pre-PCR concentrations. The remaining 25 μL reaction was placed into a preheated thermocycler and run for 8 cycles, using the following cycling conditions:

1. **94° C** for 2 min. ***Denature***
2. **94° C** for 30 sec. ***Denature***
3. **78° C** for 5 sec. ***PNA annealing***
4. **60° C** for 1 min. ***Primer annealing***
5. **72° C** for 1 min. ***Extension***
6. **GO TO STEP 2** for **7** additional cycles
7. **16° C** forever ***Hold***

Following PCR, the pre-PCR aliquots were run alongside 5 μL of post-PCR product on a 2% gel to confirm successful amplification of the reference libraries (Supplementary Figure 10). The remaining 20 μL of PCR reactions were then cleaned with SPRI beads (1.5 : 1.0 [v/v]) and set aside. A large aliquot of the original library (approximately 50 ng) was also run on a 2% gel to separate the host and microbe bands for individual purification. The bands were cut out of the gel and each band was put into a separate Econospin spin column (Epoch Life Sciences, Missouri City, TX, USA) without any other liquids or binding buffer. The gel slices were centrifuged at maximum speed to force the liquid containing the DNA into the bottom chamber, leaving the dried gel on top. The eluted DNA was cleaned with SPRI beads at 1.5 : 1.0 (v/v) and eluted in EB.

The purified pooled host library fraction, pooled microbe library fraction, and each of the four reference libraries were quantified with Picogreen and the molarity of each was estimated. The pools were then mixed together, targeting host molarity at 5% of the total and each reference library at 1% of the total.

### Illumina sequencing

Pooled and quality-checked sequencing libraries were cleaned of all remaining dimers and off-target fragments using a BluePippin (Sage Science, Beverly, MA, USA) set to a broad range of 280 to 720 bp. The libraries were then diluted for Illumina sequencing following manufacturers’ protocols. Libraries were first diluted to 2.5 - 2.8 nM in elution buffer (EB, 10 mM Tris pH 8.0) and spiked into a compatible lane of the HiSeq3000 instrument (2 × 150 bp paired end reads) to occupy 2-3% of the lane. Samples were sequenced across four total lanes (Supplementary Table 1)

### Sequence processing

The sequences were demultiplexed first by the 9 bp barcode on the PCR primers (Supplementary Table 1), of which there are 96, not allowing for any mismatches. In some cases in which two samples differed in both their host and microbe primer sets, we amplified both samples with the same 9 bp barcode to increase multiplexing; such samples were further demultiplexed using regular expressions for the forward primer and reverse primer sequences. Following demultiplexing, all samples were filtered to remove sequences with any mismatches to the expected primers. With HiSeq3000 150 bp read lengths, overlap of read 1 and read 2 was not possible for our amplicons, and therefore only read 1 was processed further.

All primer sequences were removed. Additional quality filtering, removal of chimeric sequences, preparation and Amplicon Sequence Variant (ASV) tables, and taxonomic assignment were done with a combination of VSEARCH^71^ and USEARCH10^39^. ASVs were prepared as ‘zero-radius OTUs’ (zOTUs)^39^. The 16S rDNA taxonomy was classified based on the RDP training set v16 (13k seqs.)^72^, and ITS1 taxonomy of the top 10 most abundant fungal ASVs was classified manually using the UNITE database^73^ (https://unite.ut.ee/). To reduce memory usage, data from the five lanes was processed into four independent ASV tables (Supplementary Data), as described in the sample metadata (Supplementary Table 1).

ASV tables were analyzed statistically and graphically using custom scripts in R^74^, particularly with the help of packages “ggplot2”^75^ and “reshape2”^76^. Custom scripts are available on GitHub at (https://github.com/derekLS1/hamPCR).

### Samples

#### Synthetic titration panel

Seeds from the *Arabidopsis thaliana* accession Col-0 were surface sterilized by immersion for 1 minute in 70% ethanol with 0.1% Triton X-100, soaking in 10% household bleach for 12 minutes, and washing three times with sterile water. Seeds were germinated axenically on ½ strength MS media with MES, and about 2 g of seedlings were harvested after 10 days. DNA was extracted in the sterile hood as in^12^ and diluted to 10 ng/μL in elution buffer (10 mM Tris pH 8.0, hereafter EB). Pure *E. coli* and *Sphingomonas* sp. cultures were likewise grown with LB liquid and solid media respectively, and DNA was extracted using a bead beating protocol^12^. *E. coli* DNA was used separately, or alternatively pooled with the mixed *Sphingomonas* DNA, and diluted to 10 ng/μL. The plant DNA and microbial DNA were then combined according to the following table:

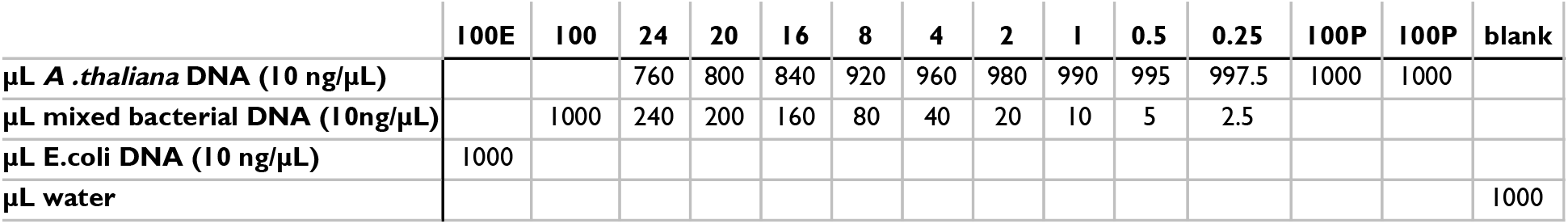

V4 tagging was performed with 515_F1_G-46603 and 799_R1_G-46601 (V4 16S rDNA) and At.GI_F1_G-46602 and At.GI_R502bp_G-46614 (*A. thaliana GI*). Each exponential PCR reaction was completed in a single reaction of 25 μL.

#### Synthetic equimolar plasmid template

The ITS1 region from *Agaricus bisporus*, a fragment of the *GI* gene from *A. thaliana* Col-0 accession, the 16S rRNA gene from *Pto* DC3000, and the ITS1 region from *H. arabidopsidis* were PCR amplified individually, combined into one fragment via overlap extension PCR, and cloned into pGEM®-T Easy (Promega, Madison, WI, USA). The sequences of these templates can be found in Supplementary Information.

#### Wild *A. thaliana* samples

DNA from chosen samples previously analyzed by conventional 16S rDNA-only sequencing of the V4 region and shotgun metagenomics^12^ was reused, chosen to capture a wide range of realistic bacterial loads. The samples were individually assayed with hamPCR using 5 μL DNA template (approximately 50 ng). V4 tagging was performed with 515_F1_G-46603 and 806_R1_G-46631 (V4 16S rDNA) and At.GI_F1_G-46602 and At.GI_R502bp_G-46614 (502 bp *A. thaliana GI*). These were the only V4 samples tagged with the 806R primer instead of the nearby 799R primer, and it was used to enable direct comparison to the dataset in^12^. V3V4 tagging was performed with 341_F1_G-46605 and 799_R1_G-46601 (V3V4 16S rDNA) and At.GI_F1_G-46602 and At.GI_R502bp_G-46614 (502 bp *A. thaliana GI*). V5V6V7 tagging was performed with 799_F1_G-46628 and 1192_R1_G-46629 (V5V6V7 16S rDNA) and At.GI_F1_G-46602 and At.GI_R466bp_G-46652 (466 bp *A. thaliana GI*). Each exponential PCR reaction was completed in a single reaction of 25 μL; each sample was replicated three times.

#### Wild *A. thaliana* mixed sample

DNA from samples previously analyzed by conventional 16S rDNA-only sequencing of the V4 region and shotgun metagenomics^12^ were pooled to prepare a single abundant mixed sample to be used repeatedly for technical tests.

#### *Hyaloperonospora arabidopsidis* and *Pseudomonas syringae pv. tomato* DC3000 co-infection

Both wildtype *A. thaliana* seedlings in the Col-0 genetic background and *enhanced disease susceptibility 1* mutants in the Ws-0 genetic background (*eds1-1)* were grown from surface-sterilized seeds. Seedlings were raised in ED73 potting mix (Einheitserdewerke, Sinntal-Altengronau, Germany) in 5 cm pots for 10 days under short-day conditions (8 hours light, 16 hours dark). Each pot contained 4 to 5 seedlings, and for each genotype, four pots were used for each infection condition. Plants were treated with either 10 mM MgCl_2_ (buffer only), *H. arabidopsidis* (*Hpa*) isolate 466-1 alone (5 × 10^4^ spores / mL), or *Hpa* 466-1 with *P. syringae pv. tomato* (*Pto*) DC3000 (OD_600_ = 0.25, a gift from El Kasmi lab, University of Tübingen).

The infected plants were grown at 16°C for 8 days (10 hours light, 14 hours dark) and harvested by pooling all seedlings in each pot into a sterile pre-weighed tube, which was again weighed to find the mass of the seedlings. Three 5 mm glass balls and 300 μL 10 mM MgCl_2_ were added to each tube and the plant cells were lysed at a speed of 4.0 m/s for 20 seconds in a FastPrep-24™ Instrument (MP Biomedicals, Illkirch-Graffenstaden, France) to release the live bacteria from the leaves. From the pure lysate, 20 μL was used for a serial log dilution series, and 5 μL of each dilution was plated on LB agar supplemented with 100 μg/mL rifampicin. Colony forming units (CFUs) were counted after 2 days of incubation at 28°C. The remaining 280 μL of lysate were combined with 520 μL DNA lysis buffer, 0.5 mL of 1 mm garnet sharp particles (BioSpec, Bartlesville, OK, USA). 60 μL of 20% SDS was added to make a final SDS concentration of 1.5%, and DNA was extracted using a bead beating protocol^12^. The number of *Hpa* sporangiophores was too high to be accurately quantified by visual counting.

The DNA preps were individually assayed with hamPCR using 5 μL DNA template (approximately 30 ng); tagging was performed with three primer sets: Ha.Actin_F1_G-46716 and Ha.Actin_R1_G-46717 (Hpa Actin), At.GI_F1_G-46602 and At.GI_R502bp_G-46614 (502 bp *A. thaliana GI*), and 515_F1_G-46603 and 799_R1_G-46601 (V4 16S rDNA). Each exponential PCR reaction was completed in three parallel reactions of 13 μL, which were recombined prior to sequencing.

#### Titration with plant DNA infected with *H. arabidopsidis*

A titration panel was made combining different amounts of DNA from uninfected plants (*eds1-1* treated only with 10 mM MgCl_2_) and DNA from *Hpa*-infected plants (*eds1-1* infected with *Hpa* as described above). Infected and uninfected pools were each diluted to 6 ng/μL, and combined in 0:7, 1:6, 2:5, 3:4, 4:3, 5:2, 6:1, and 7:0 ratios. These were tagged using the same three primer sets described above for *Hpa* actin, *A. thaliana GI*, and V4 16S rDNA above. Each exponential PCR reaction was completed in a single reaction of 25 μL; hamPCR was replicated on the titration three times.

#### *Capsicum annuum* infections with *Xanthomonas*

##### Leaf infiltration log series

Using pressure infiltration with a blunt-end syringe, *C. annuum* cultivar Early Calwonder (ECW) leaves were inoculated with *Xanthomonas euvesicatoria* (*Xe*). *Xe* strain 85-10 ^53^ was resuspended in 10 mM MgCl_2_ to final concentration of 10^8^ CFU / mL (OD_600_=0.4) and further diluted to 10^7^, 10^6^, 10^5^ and 10^4^ CFU / mL. Upon infiltration, 5 leaf discs (7 mm diameter) were punched from each leaf per sample and placed in a 2 mL round-bottom tube with two SiLibeads (type ZY-S 2.7-3.3 mm, Sigmund Lindner GmbH, Warmensteinach, Germany) and 300 μL 10 mM MgCl_2_. The samples were ground by bead beating for 25 sec at 25 Hz using a Tissue Lyser II machine (Qiagen, Hilden, Germany). For CFU-based bacterial enumeration, 30 μL of the lysate or 30 μL of serial dilutions were plated on NYG medium (0.5 % peptone, 0.3 % yeast extract, 0.2 % glycerol and 1.5 % agar ^77^ containing rifampicin (100 μg/ml). *Xe* bacteria were counted 3 days post incubation at 28°C. The remaining 250 μL of lysate was combined with 600 μL of DNA lysis buffer containing 2.1% SDS (for a 1.5% final SDS concentration) and transferred to screw cap tubes filled with 1 mm garnet sharp particles, for a bead-beating DNA prep as previously described^12^.

##### Growth curve

*Xe* strain 85-10, resuspended in 10 mM MgCl_2_ to a final concentration of 10^4^ CFU / mL, was infiltrated via a blunt end syringe into 6 *C. annuum* (ECW) leaves of 6 different plants. Upon 0, 2, 4, 7, 9 and 11 dpi (days post inoculation) 4 leaf discs (7 mm diameter) from each inoculated leaf were harvested and bacterial numbers were determined as described above. 250 μL of leaf lysates were used for a bead-beating DNA prep as described for all other samples above.

Each hamPCR template tagging reaction used 5-10 μL template (approximately 50 ng each); tagging was performed with primers 515_F3_G-46694 and 799_R1_G-46601 (V4 16S rDNA), and Ca.GI_F1_G-46626 and Ca.GI_R1_G-46627 (*C. annuum GI*). Each exponential PCR reaction was completed in three parallel reactions of 13 μL, which were recombined prior to sequencing.

#### *Pristionchus pacificus* titration panel

*Pristionchus pacificus* strain PS312^78^ was grown on nematode growth media (NGM) plates supporting a bacterial lawn of either pure *E. coli* OP50 or alternatively a mix of *E. coli* OP50, *Pto* DC3000, and *Lysinibacillus xylanilyticus* (a strain isolated from wild *P. pacificus*). The worms were washed extensively with PBS buffer to remove epidermally-attached bacteria, and DNA was prepared from whole worms using the same bead beating protocol as described for *A. thaliana* ^12^. Worm DNA from the pure culture and the mixed culture were each diluted to 6 ng/μL, and combined in 0:7, 1:6, 2:5, 3:4, 4:3, 5:2, 6:1, and 7:0 ratios to create a titration panel. Each hamPCR template (5 μL template or 30 ng total) was used to perform the tagging reaction, using primers 799F1_G-46628 and 1192R1_G-46629 (V5V6V7 16S rDNA), and Pp_csq-1_F1_G-46691 and Pp_csq-1_R1_G-46692 (*P. pacificus csq-1*). Each exponential PCR reaction was completed in a single reaction of 25 μL; the titration was replicated three times.

#### *Triticum aestivum* titration panel

*Triticum aestivum* (wheat) seeds (Rapunzel Naturkost, Legau, Germany) were surface-sterilized by immersion in 70% ethanol and 0.1% Triton X-100 for 1 minute, soaking for 15 minutes in 10% household bleach, and finally washing three times in sterile autoclaved water. Axenic plants were grown on 1% agar supplemented with 1/2 strength MS medium buffered with MES. About 1 g of sterile leaf tissue was harvested after 10 days, and DNA was extracted in the sterile hood as described in ref.^12^. Roots that had been spontaneously colonized by microbes were obtained by growing by transplanting germinated seeds outdoors into potting soil. Roots were harvested from approximately 4-week old plants and surface-sterilized by immersion in 10% household bleach with 0.1% Triton X-100 for 5 minutes, followed by 3 washes with sterile water. Axenic leaf DNA and spontaneously-colonized root DNA were each diluted to 60 ng/μL and combined in 0:7, 1:6, 2:5, 3:4, 4:3, 5:2, 6:1, and 7:0 ratios to create a titration panel of eight samples. Each hamPCR tagging reaction used 3 μL (~180 ng) template; fungal ITS1 tagging was performed with primers ITS1_F1_G-46622 and ITS2_R1_G-46623 (ITS1 rDNA), and PolA1_F1_G-46750 and PolA1_R1_G-46751 (*T. aestivum* RNA polymerase 1 gene, *PolA1*). Bacterial 16S rDNA tagging was performed with the same *PolA1* primers and with 515_F1_G-46603 and 799_R1_G-46601 (V4 16S rDNA). To make an additional ITS1 library enriched for ITS1 amplicons, the ITS1 primer pair concentration was increased by a factor of 1.33 and the *PolA1* primer pair concentration was decreased by a factor of 0.66, giving a 2:1 ratio instead of the standard 1:1 ratio, and the tagged products were amplified with 7 tagging and 25 PCR cycles instead of the standard 2 tagging and 30 PCR cycles.

#### *Zea mays* field samples

Samples of leaves from mature *Zea mays* (maize) genotype B73 were harvested by standard hole punch from a field side in Tübingen. Permission to punch the leaves was graciously provided by Dr. Marja Timmermans (University of Tübingen). Each sample comprised 5 leaf discs, which were immediately shaken in 1 mL of sterile water in a screw cap tube to remove dust from the field. The water was removed by pipetting and the leaf discs were snap frozen in liquid nitrogen and taken back to the lab for processing. DNA was extracted with the bead beating protocol described above, with the difference that prior to addition of lysis buffer and garnet rocks, the deep frozen leaf discs were pre-ground with 3 metal ball bearings at a speed of 5.0 m/s in a FastPrep-24™ instrument. We found this pre-grind was helpful to break down the fibrous maize leaf tissue. Prior to adding garnet rocks and lysis buffer, the metal balls were removed by magnet, as metal balls can crack the tubes at the speed of 6.0 m/s used for the primary DNA extraction. Each hamPCR tagging reaction used 10 μL (~120 ng) template; Bacterial 16S rDNA was tagged with one of the forward primers 515F_bcGA_G-47188, 515F_bcTC_G-47189, 515F_bcAG_G-47190, or 515_F3_G-46694 paired with the reverse primer 799_R1_G-46601 (V4 16S rDNA). Maize *LUMINIDEPENDENS (LD)* was tagged with one of the forward *LDP1* primers Zm_LD_bcGA_G-47184, Zm_LD_bcTC_G-47185, Zm_LD_bcAG_G-47186, or Zm_LD_bcCT_G-47187 paired with the *LD* reverse primer Zm_LD_R_G-47158. the standard 2 tagging and 30 PCR cycles. Two tagging cycles were paired with 30 exponential cycles. To reduce host representation in the final library from the original ~75% to approximately 40%, we used the gel remixing technique described in Figure 3.

### Test of tagging step cycle numbers

As templates, we used a pool of mixed wild *A. thaliana* leaf DNA (~ 50 ng / reaction) and the “synthetic equimolar plasmid template” (~ 0.05 ng / reaction). For the wild *A. thaliana* leaf DNA, we tested V4 16S rDNA primers alone in the tagging step vs. hamPCR with V4 16S rDNA primers plus primers for the host *GI* gene. For the “synthetic equimolar plasmid template”, we used only hamPCR. Specifically, we used 515_F1_G-46603 and 799_R1_G-46601 (V4 16S rDNA) and At.GI_F1_G-46602 and At.GI_R502bp_G-46614 (502 bp *GI* gene).

We applied hamPCR for 2, 3, 4, 5, 6, 7, 8, 9, or 10 tagging cycles, paired with 30, 29, 28, 27, 26, 25, 24, 23, or 22 PCR cycles, respectively. All tagging and PCR reactions were started together, and fewer tagging cycles than 10, or fewer PCR cycles than 30, were achieved by taking PCR tubes out of the thermocycler at the end of the appropriate extension steps and placing them on ice.

### Tests of template concentrations

A panel of 8 concentrations of wild *A. thaliana* leaf DNA was prepared, ranging from 5 to 500 ng per reaction. Primers for the wild *A. thaliana* leaf DNA were 515_F1_G-46603 and 799_R1_G-46601 (V4 16S rDNA) and At.GI_F1_G-46602 and At.GI_R502bp_G-46614 (502 bp *GIGANTEA* gene), with both primer pairs in equal ratio.

### Quantitative real time PCR on *C. annuum* samples

A primer set targeting the gene for the type III effector XopQ of pathogenic *Xanthomonas* was used to measure abundance of *Xe* 85-10^54^. For *C. annuum*, primers targeting the *UBI-3* gene encoding a ubiquitin-conjugating protein were used ^55^. Two reagent mastermixes were prepared, one for each primer set, to help improve primer dose consistency. Each sample was amplified using three 10 μL technical replicates per primer set that were averaged for analysis. Each 10 μL reaction included 2.5 μL of DNA, to which was added, as a mastermix, 5 μL SYBR® Green PCR Master Mix (Life Technologies, Carlsbad, California), 1.5 μL water, 0.5 μL of 5 μM forward primer, and 0.5 μL of 5 μM reverse primer. qPCR was performed on a BioRad CFX384 Real-time System and analyzed with the CFX Manager Software. The following conditions were used for the amplification of both target genes:

1. **94° C** for 5 min. ***Denature***
2. **94° C** for 30 sec. ***Denature***
3. **55° C** for 30 sec. ***Annealing***
4. **68° C** for 45 sec. ***Extension***
5. **Image fluorescence**
6. GO TO STEP **2** for **39** additional cycles

The ratio of microbial to host DNA was initially calculated as 2^(-mean *xopQ* Cq value) / 2^(-mean *UBI-3* Cq value). See alignment to CFU counts below.

### Alignment of Xanthomonas qPCR and hamPCR load with CFU counts

Log10-transformed *Xe* 85-10 ASV loads from hamPCR were regressed onto log10-transformed *xopQ* loads from qPCR (least squares method), and the slope (*m*) and y-intercept (*b*) of the best-fit line were used to transform and align the qPCR loads to hamPCR loads with the following formula: Load_qPCR_hamPCR-aligned_ = *m* × Load_qPCR + *b*. Next, log10-transformed CFU counts were regressed onto the log10-transformed hamPCR loads, and the slope and y-intercept of the resulting best-fit line were used similarly to align both Load_qPCR_hamPCR-aligned_ and hamPCR loads to the CFU counts.

### Correlation networks

Pearson correlation matrices for relative abundance and microbial load data were created in R^74^ using the “stats” package. The SparCC^56^ correlation matrix was created in R using the implementation in the “SpiecEasi” package^79^. Networks were visualized with the package “qgraph”^80^. Custom scripts are available on GitHub (https://github.com/derekLS1/hamPCR).

## Supporting information

R168_ASV_table

R175L1_ASV_table

R175L4_ASV_table

R177L5_ASV_table

## Data availability

All data in this manuscript has been deposited in the European Nucleotide Archive (ENA). It can be accessed under the project number PRJEB38287. At https://www.ebi.ac.uk/ena.

## Author contributions

DL and PP planned the study. DL led the data analysis. DL and PP performed all DNA extractions and prepared all hamPCR libraries for sequencing. PP constructed the synthetic plasmid template. AS and TL planned, and AS conducted, the pepper and *Xanthomonas* infections. DL and GS planned and conducted the *Hpa* and *Pto* DC3000 co-infection of *A. thaliana*. WL grew and harvested the colonized *P. pacificus* samples. DL, PP, and DW wrote the manuscript.

## Acknowledgements

We thank Lei Li and Fiona Beitel for discussions and suggestions regarding PCR and qPCR, Talia Karasov for suggestions on presentation, Katrin Fritschi and Heike Budde for technical help with sequencing, Ilja Bezrukov for assistance with demultiplexing, and Dor Russ, Jeff Dangl, and Benjamin Schwessinger for supportive discussions and comments on the manuscript. Thanks to Victor Schmidt, Nick Youngblut, Lei Li, and Hua Wang for giving us space in their HiSeq 3000 lanes. Thanks to Marja Timmermans for access to field maize plants. Supported by a Human Frontiers Science Program (HFSP) Long-Term Fellowship (LT000565/2015-L to DL), the DFG through Priority Program SPP DECRyPT, and the Max Planck Society (DW).

