## Supplementary_Information for "Host-associated microbe PCR (hamPCR): accessing new biology through convenient measurement of both microbial load and community composition"


Supplementary Figure 1 2
Primers: Microbe Tagging 4
- V4 region (16S rDNA), 515F - 799R / 515F - 806R, ~414 bp final amplicon 4
- V3V4 region (16S rDNA), 341F - 799R, ~587 bp final amplicon 4
- V5V6V7 (16S rDNA) region, 799F - 1192R, ~543 bp final amplicon 5
- ITS oomycete, ITS1-O_F - 5.8s-O_R, ~394 bp final amplicon 5
- ITS1 fungus, ITS1F - ITS2, variable length final amplicon 6
- Actin gene from *Hyaloperonospora arabidopsidis*, 585 bp final amplicon 6

Primers: Host Tagging 7
- *GIGANTEA (GI)* from *Arabidopsis thaliana*, 502 bp final amplicon 7
- *GIGANTEA (GI) from Arabidopsis thaliana*, 466 bp final amplicon 7
- *GIGANTEA (GI)* from *Capsicum annuum*, 508 bp final amplicon 8
- RNA polymerase I gene (*PolA1*) from *Triticum aestivum*, 497 bp final amplicon 8
- Calsequestrin gene (*csq-1*) from *Pristionchus pacificus*, 470 bp final amplicon 8
- *LUMINIDEPENDENS (LD)* from *Zea mays*, 470 bp final amplicon 9

Note on Adapting Tagging Sets for Other Protocols 10
Primers: Exponential PCR 11
Supplementary Figure 2 12
Supplementary Figure 3 13
Supplementary Figure 4 14
Supplementary Figure 5 15
Supplementary Figure 6 16
Supplementary Figure 7 17
Supplementary Figure 8 18
Supplementary Figure 9 19
Supplementary Figure 10 20
Supplementary Figure 11 21
Supplementary Figure 12 22
Supplementary Figure 13 24
Supplementary Figure 14 25
Supplementary Figure 15 26
Supplementary Figure 16 28
Supplementary Figure 17 29
Supplementary Discussion: Synthetic Template 30
Supplementary References 32

### Supplementary Figure 1

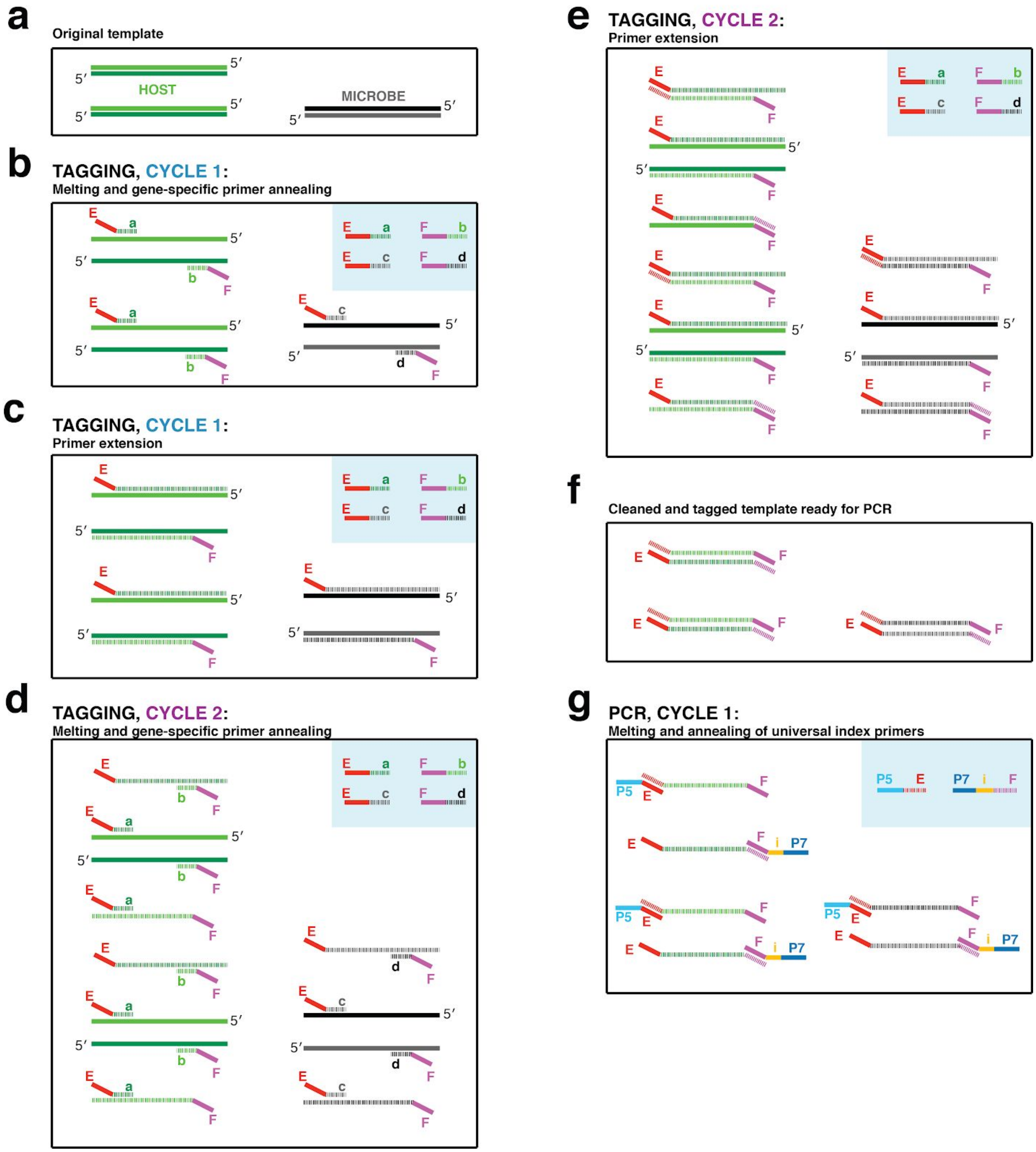


**Supplementary Figure 1. PCR Scheme | a**, The input DNA is shown as two double-stranded DNA molecules from the host (light/dark green) and one double-stranded DNA molecule from a microbe (black/grey). **b**, In the annealing step of tagging cycle one, an excess of two primer pairs is present (blue box), each with a gene-specific region (regions a through d) and universal overhangs (regions E and F). The gene-specific regions of each primer anneal to the templates. **c**, In the extension step of tagging cycle 1, primers extend to make a single copy of each template molecule with a universal overhang on the 5' end, producing "single-tagged templates". Extended molecules are represented by dashed lines, while original templates retain a solid line. **d**, In tagging cycle 2, again the gene-specific regions of each primer anneal to all template molecules, both to the originals and single-tagged templates. **e**, In the extension step, note that for those primers that had annealed to single-tagged templates, extension generates the reverse complement of the universal overhang, producing "double-tagged templates". **f**, Primers are removed from the reaction with SPRI beads prior to PCR. Note that the quantity of double-tagged template molecules is the same as the quantity of original template molecules. Although original template and single-tagged templates survive SPRI cleanup and are also present in this step, these lack the universal overhangs and cannot be amplified in PCR and are not shown. **g**, For the exponential PCR step, an excess of a single primer pair is present (blue box), each with a region complementary to the universal overhangs (regions E and F) and sequencing adapters (regions P5 and P7). An index for multiplexing is present on one or both primers (region i). Because double-tagged templates from both host and microbe have the same universal overhangs, they are not differentially amplified during PCR.

### Primers: Microbe Tagging

Standard desalting is sufficient purification for the tagging primers.

#### V4 region (16S rDNA), 515F - 799R / 515F - 806R, ~414 bp final amplicon

*Underlined letters represent the universal overhang.*
*799F is commonly used in combination with 1192R because it avoids plant plastids by mismatch. Shown here is the reverse complement of that primer. Used in combination with PNAs (Lundberg et al., 2013), it avoids nearly all plastids.*
*Bracketed grey boxed letters represent "linkers" - sequences designed to anneal to as few 16S rDNA molecules as possible, and ensure that only the gene-specific region binds to the template.*
*Lowercase letters are template-specific. Template-specific primer sequences for 515F and 806R are the same as used in the Earth Microbiome Project (Thompson et al., 2017) and (Lundberg et al., 2013). Linker sequences for 515F and 806R are as in (Lundberg et al., 2013). Template-specific primer sequence for 799F as in (Agler et al., 2016).*
*"F1", "F3", "bcGA", "bcTC", and "bcAG" forward primers perform identically but each introduces different sequenced bases which serve as an additional barcode. See (Lundberg et al., 2013).*

**FORWARD:**
>515_F1_G-46603
TCCCTACACGACGCTCTTCCGATCT [GA] **gtgycagcmgccgcggtaa**
>515_F3_G-46694
TCCCTACACGACGCTCTTCCGATCT CT [GA] **gtgycagcmgccgcggtaa**
>515F_bcGA_G-47188
TCCCTACACGACGCTCTTCCGATCT GA [GA] **gtgycagcmgccgcggtaa**
>515F_bcTC_G-47189
TCCCTACACGACGCTCTTCCGATCT TC [GA] **gtgycagcmgccgcggtaa**
>515F_bcAG_G-47190
TCCCTACACGACGCTCTTCCGATCT AG [GA] **gtgycagcmgccgcggtaa**

**REVERSE 799:**
>799_R1_G-46601
GGAGTTCAGACGTGTGCTCTTCCGATCT [TG] **cmgggtatctaatcckgtt**

**REVERSE 806:**
>806_R1_G-46631
GGAGTTCAGACGTGTGCTCTTCCGATCT [AC] **ggactacnvgggtwtctaat**

>E.coli_V4_sequenced_region_285bp_final_amplicon_413bp_
**gtgccagcagccgcggtaa**TACGGAGGGTGCAAGCGTTAATCGGAATTACTGGGCGTAAAGCGCACGCAGGCG
GTTTGTTAAGTCAGATGTGAAATCCCCGGGCTCAACCTGGGAACTGCATCTGATACTGGCAAGCTTGAGTCTC
GTAGAGGGGGGTAGAATTCCAGGTGTAGCGGTGAAATGCGTAGAGATCTGGAGGAATACCGGTGGCGAAGGCG
GCCCCCTGGACGAAGACTGACGCTCAGGTGCGAAAGCGTGGGGAGCA**aacaggattagataccctg**

#### V3V4 region (16S rDNA), 341F - 799R, ~587 bp final amplicon

*Underlined letters represent the universal overhang.*
*799F is commonly used in combination with 1192R because it avoids plant plastids by mismatch. Shown here is the reverse complement of that primer. Used in combination with PNAs (Lundberg et al., 2013), it avoids nearly all plastids.*
*Bracketed grey boxed letters represent "linkers" - sequences designed to anneal to as few 16S rDNA molecules as possible, and ensure that only the gene-specific region binds to the template.*

*Lowercase letters are template-specific. Template-specific primer sequence for 341F is as in (Agler et al., 2016).*

**FORWARD:**
>341_F1_G-46605
TCCCTACACGACGCTCTTCCGATCT **[GA]** **cctacgggaggcagcag**
**REVERSE:**
>799_R1_G-46601
GGAGTTCAGACGTGTGCTCTTCCGATCT **[TG]** **cmgggtatctaatcckgtt**

>E.coli_V3V4_sequenced_region_459bp_final_amplicon_587bp
**cctacgggaggcagcag**TGGGGAATATTGCACAATGGGCGCAAGCCTGATGCAGCCATGCNGCGTGTATGAAG
AAGGCCTTCGGGTTGTAAAGTACTTTCAGCGGGGAGGAAGGGAGTAAAGTTAATACCTTTGCTCATTGACGTT
ACCCGCAGAAGAAGCACCGGCTAACTCCGTGCCAGCAGCCGCGGTAATACGGAGGGTGCAAGCGTTAATCGGA
ATTACTGGGCGTAAAGCGCACGCAGGCGGTTTGTTAAGTCAGATGTGAAATCCCCGGGCTCAACCTGGGAACT
GCATCTGATACTGGCAAGCTTGAGTCTCGTAGAGGGGGGTAGAATTCCAGGTGTAGCGGTGAAATGCGTAGAG
ATCTGGAGGAATACCGGTGGCGAAGGCGGCCCCCTGGACGAAGACTGACGCTCAGGTGCGAAAGCGTGGGGAG
CA**aacaggattagataccctg**

### V5V6V7 (16S rDNA) region, 799F - 1192R, ~543 bp final amplicon

*Underlined letters represent the universal overhang.*
*Bracketed grey boxed letters represent "linkers" - sequences designed to anneal to as few 16S rDNA molecules as possible, and ensure that only the gene-specific region binds to the template.*
*Lowercase letters are template-specific. Template-specific primer sequences are as in (Agler et al., 2016).*

**FORWARD:**
>799 F1 G-46628
TCCCTACACGACGCTCTTCCGATCT **[GT]** **aacmggattagataccckg**

**REVERSE:**
>1192 R1 G-46629
GGAGTTCAGACGTGTGCTCTTCCGATCT **[GT]** **acgtcatccccaccttcc**

>E.coli V5V6V7 sequenced region 414bp final amplicon 543bp
**aacaggattagataccctg**GTAGTCCACGCCGTAAACGATGTCGACTTGGAGGTTGTGCCCTTGAGGCGTGGC
TTCCGGAGCTAACGCGTTAAGTCGACCGCCTGGGGAGTACGGCCGCAAGGTTAAAACTCAAATGAATTGACGG
GGGCCCGCACAAGCGGTGGAGCATGTGGTTTAATTCGATGCAACGCGAAGAACCTTACCTGGTCTTGACATCC
ACGGAAGTTTTCAGAGATGAGAATGTGCCTTCGGGAACCGTGAGACAGGTGCTGCATGGCTGTCGTCAGCTCG
TGTTGTGAAATGTTGGGTTAAGTCCCGCAACGAGCGCAACCCTTATCCTTTGTTGCCAGCGGTCCGGCCGGGA
ACTCAAAGGAGACTGCCAGTGATAAACTGGA**ggaaggtggggatgacgt**

### ITS oomycete, ITS1-O_F - 5.8s-O_R, ~394 bp final amplicon

*Underlined letters represent the universal overhang.*
*Lowercase letters are template-specific. Template-specific primer sequences are as in (Agler et al., 2016)*

**FORWARD:**
>ITS1-O_F1_G-46636
TCCCTACACGACGCTCTTCCGATCT **cggaaggatcattaccac**

**REVERSE:**
>5.8s-O_R1_G-46637
GGAGTTCAGACGTGTGCTCTTCCGATCT **agcctagacatccactgctg**

>ITS_H.arabidopsidis_sequenced_region_269bp_final_amplicon_394bp
**cggaaggatcattaccac**ACCTAAAAAACTTTCCACGTGAACCGTTTCAACCCAATAGTTGGGGGTCTTATTT
GGCGGCGGCTGCTGGCTTAATTGTTGGCGGCTGCTGCTGAGTGAGCCCTATCAAAAAAAAGGCGAACGTTTGG
GCTTCGGCCTGATTTAGTAGTCTTTTTTTCTTTTAAACCCCTTCCTTAATACTGAATATACTGTGGGGACGAA
AGTCTCTGCTTTTAACTAGATAGCAACTTT**cagcagtggatgtctaggct**

### ITS1 fungus, ITS1F - ITS2, variable length final amplicon

*Underlined letters represent the universal overhang.*
*Bracketed grey boxed letters represent "linkers" - sequences designed to anneal to as few ITS molecules as possible, and ensure that only the gene-specific region binds to the template.*
*Lowercase letters are template-specific. Template-specific primer sequences and linker sequences are as used in the Earth Microbiome Project (Thompson et al., 2017).*

**FORWARD:**
>ITS1_F1_G-46622
TCCCTACACGACGCTCTTCCGATCT **[GG]** **cttggtcatttagaggaagtaa**

**REVERSE:**
>ITS2_R1_G-46623
GGAGTTCAGACGTGTGCTCTTCCGATCT **[CG]** **gctgcgttcttcatcgatgc**

>ITS1__Agaricus_bisporus_sequenced_region_407bp_final_amplicon_532bp
**cttggtcatttagaggaagtaa**AAGTCGTAACAAGGTTTCCGTAGGTGAACCTGCGGAAGGATCATTATTGAA
TTATGTTTTCTAGATGGGTTGTAGCTGGCTCTTCGGAGTATGTGCACGCCTGTCTGGACTTCATTTTCATCCA
CCTGTGCACCTTTTGTAGTCTTTTTCAGGTATTGGAGGAAGTGGTCAGCCTATCAGCTCTTTGCTGGATGTAA
GGACTTGCAGTGTGAAAACAGTGCTGTCCTTTACCTTGGCCATGGAATCTTTTTCCTGTTAGAGTCTATGTTA
TTCATTATACTCTTAGAATGTCATTGAATGTCTTTACATGGGCTATGCCTATGAAAATTATTATACAACTTTC
AGCAACGGATCTCTTGGCTCTC**gcatcgatgaagaacgcagc**

### Actin gene from *Hyaloperonospora arabidopsidis*, 585 bp final amplicon

*Underlined letters represent the universal overhang.*
*Lowercase letters are template-specific. The template-specific primer sequences were chosen for this study; the choice of the Actin gene was based on (Anderson & McDowell, 2015).*

>Ha.Actin_F1_G-46716
TCCCTACACGACGCTCTTCCGATCT **cgcgctgccgcacgcgattgt**

>Ha.Actin_R1_G-46717
GGAGTTCAGACGTGTGCTCTTCCGATCT **cgccaaagccgtcagttccttc**

>Actin_Hyaloperonospora_arabidopsidis_Emoy2_sequenced_region_457bp_final_
amplcion_585bp
**cgcgctgccgcacgcgattgt**GCGTTTGGATCTCGCTGGTCGCGACTTGACCGACTACATGATGAAGATCTTG
ACGGAGCGCGGGTACTCGTTTACTACCACGGCCGAGCGCGAAATCGTGCGCGACATTAAAGAGAAACTCACGT
ACATTGCACTGGACTTTGACCAGGAGATGAAGACAGCGGCCGAGTCGTCGGGACTCGAGAAGAGCTACGAATT
GCCGGATGGCAATGTGATTGTCATTGGCAATGAACGTTTCCGTACGCCGGAAGTGCTGTTCCAGCCGTCGCTC
ATTGGCAAAGAAGCTGCCGGTATTCACGACTGCACGTTCCAGACCATCATGAAGTGTGACGTGGATATCCGGA
AGGATTTGTACTGCAACATTGTGCTCTCGGGCGGAACCACCATGTACCCGGGCATTGGCGAACGCATGAC**gaa
ggaactgacggctttggcg**

### Primers: Host Tagging

#### *GIGANTEA* (*GI*) from *Arabidopsis thaliana*, 502 bp final amplicon

(Size distinguishable from V4 or V3V4 16S rDNA primers)

*Underlined letters represent the universal overhang.*

*Lowercase letters are template-specific. Here, the template-specific portion extends into the universal overhang for the At.GI_F1_G-46602 primer.*

*"F1" and "F3" forward primers perform identically but F3 introduces a "CT" causing a frameshift that can also serve as an additional barcode. See (Lundberg et al., 2013). Here, the additional "CT" extends into the universal overhang for the At.GI_F3_G-46640 primer.*

**FORWARD:**

>At.GI_F1_G-46602
TCCCTACACGACGCTCTTCCGAT**ct gtaaagataaatgggtcatctaa**
>At.GI_F3_G-46640
TCCCTACACGACGCTCTTCCGAT CT **ctgtaaagataaatgggtcatctaa**

**REVERSE:**

>At.GI_R502bp_G-46614
GGAGTTCAGACGTGTGCTCTTCCGATCT **tccttctgaaccggtgtattc**

>Athaliana_GI_sequenced_region_377bp_final_amplicon_502bp
**gtaaagataaatgggtcatctaa**AGAGTATGGAGCTGGGATTGACTCGGCAATTAGTCATACGCGCCGAATTT
TGGCAATCCTAGAGGCACTCTTTTCATTAAAACCATCTTCTGTGGGGACTCCATGGAGTTACAGTTCTAGTGA
GATAGTTGCTGCGGCCATGGTTGCAGCTCATATTTCCGAACTGTTCAGACGTTCAAAGGCCTTGACGCATGCA
TTGTCTGGGTTGATGAGATGTAAGTGGGATAAGGAAATTCATAAAAGAGCATCATCATTATATAACCTCATAG
ATGTTCACAGCAAAGTTGTTGCCTCCATTGTTGACAAAGCTGAACCCTTGGAAGCCTACCTTAA**gaatacacc
ggttcagaagga**

#### *GIGANTEA* (*GI*) from *Arabidopsis thaliana*, 466 bp final amplicon

(Size distinguishable from V5V6V7 16S rDNA primers)

*Underlined letters represent the universal overhang.*

*Lowercase letters are template-specific. Here, the template-specific portion extends into the universal overhang for the At.GI_F1_G-46602 primer.*

**FORWARD:**

>At.GI_F1_G-46602
TCCCTACACGACGCTCTTCCGAT**ct gtaaagataaatgggtcatctaa**

**REVERSE:**

>At.GI_R466bp_G-46652
GGAGTTCAGACGTGTGCTCTTCCGATCT **aagggttcagctttgtcaacaa**

>A.thaliana_GI_sequenced_region_341bp_final_amplicon_466bp
**gtaaagataaatgggtcatctaa**AGAGTATGGAGCTGGGATTGACTCGGCAATTAGTCATACGCGCCGAATTT
TGGCAATCCTAGAGGCACTCTTTTCATTAAAACCATCTTCTGTGGGGACTCCATGGAGTTACAGTTCTAGTGA
GATAGTTGCTGCGGCCATGGTTGCAGCTCATATTTCCGAACTGTTCAGACGTTCAAAGGCCTTGACGCATGCA
TTGTCTGGGTTGATGAGATGTAAGTGGGATAAGGAAATTCATAAAAGAGCATCATCATTATATAACCTCATAG
ATGTTCACAGCAAAGTTGTTGCCTCCA**ttgttgacaaagctgaaccctt**

### ***GIGANTEA* (*GI*) from *Capsicum annuum*, 508 bp final amplicon**

(Size distinguishable from V4 or V3V4 16S rDNA primers)

*Underlined letters represent the universal overhang.*
*Lowercase letters are template-specific.*

**FORWARD:**

```
>Ca.GI_F1_G-46626
TCCCTACACGACGCTCTTCCGATCT caaatgattcatctattgagcta
```

**REVERSE:**

```
>Ca.GI_R1_G-46627
GGAGTTCAGACGTGTGCTCTTCCGATCT ctcctttagaactggtgcatg
```

```
>C.annuum_GI_sequenced_region_371bp_final_amplicon_508bp
caaatgattcatctattgagctaCGAAATGGGATTCATTCTGCCGTCTCTCATACTCGGAGGATATTGGCAAT
TTTAGAGGCACTTTTTTCTCTGAAACCATCGTCTGTTGGAACCTCATGGAGCTACAGCTCAAATGAGATAGTT
GCTGCAGCTATGGTAGCTGCTCACATTTCTGATCTGTTTAGACACAACAAGGCCTGCATGCAAGCTCTTTCTA
TTTTGATACGGTGTAAGTGGGATAATGAAATTCATTCCAGGGCATCGTCACTTTATAACCTAATTGATATTCA
TAGCAAAACTGTTGCATCAATTGTCAACAAGGCTGAACCATTGGAAGCTTATCTAATAcatgcaccagttcta
aaggag
```

### **RNA polymerase I gene (*PolA1*) from *Triticum aestivum*, 497 bp final amplicon**

(size distinguishable from V4 or V3V4 16S rDNA primers)

*Underlined letters represent the universal overhang.*
*Lowercase letters are template-specific. Here, the template-specific portion extends into the universal overhang for both primers.*

**FORWARD:**

```
>Ta.PolA1 F1 G-46750
TCCCTACACGACGCTCTTCCGATct gatgttgtggaaggaattgaa
```

**REVERSE:**

```
>Ta.PolA1 R1 G-46751
GGAGTTCAGACGTGTGCTCTTCCGAtct gcatcagctcccaagtc
```

```
>T.aestivum PolA1 sequenced region 370bp final amplicon 497bp
CTGATGTTGTGGAAGGAATTGAAGTATGCACAGTTCCTTTTCACAACAGTAATGGGCATATTTCGAGTCTCTA
TAAGTTGCATCTAAAACTATTCTCACCCGACTGTTACCCTCCTGAGTCGGAACTTACAGTAGATGAGTGTCAA
GCATCCTTGAGAACTGTGTTTGTTGATGCAATGGAATATGCAATCGAAAAACACCTAAATTTGCTACACAAAG
TTAGTGGAATCCAGGAAACAAGGGTAAAGGACACTGAGAGTTTACCATCAGAAGGTCCTGAAGAATCCGAGGG
CAGACCTACCAACGGGGACGAGTCTGATACGAGTGATGGTGACGATGAAAATGAGGATGACTTGGGAGCTGAT
GCAGA
```

### **Calsequestrin gene (*csq-1*) from *Pristionchus pacificus*, 470 bp final amplicon**

(size distinguishable from V5V6V7 16S rDNA primers)

*Underlined letters represent the universal overhang.*
*Lowercase letters are template-specific.*

**FORWARD:**

```
>Pp.csq-1_F1_G-46691
TCCCTACACGACGCTCTTCCGATCT catcgagaatatatggccgat
```

**REVERSE:**
>Pp.csq-1_R1_G-46692
GGAGTTCAGACGTGTGCTCTTCCGATCT **tgagttatgccccactattcaa**

>P.pacificus_csq-1_sequenced_region_343bp_final_amplicon_470bp
**catcgagaatatatggccgat**CGAGAATGTTCCGATTTGCCAGTGGAGAGATGTACAGGATGGAATGTCACAT
GCAGATGTGCTTGAAGGCATCGACTTGCGGGCCGAGGACTACTGTAAGTTACATAGAAACCAAAGATTGACTT
TCTAGTTGGTTCAAAATTTTCTCGAAAAGGATTTGAGAATATATAACAATTGAGAACTATTCTAATTGAGTAG
CAGTTAGACTATAATTACCGAATATTCTTTGAATGCTCACTAATACTCAATTTCCCCTGTCGTTATGCTCGTA
ATTCACCATGCAGTACCTCTTCCTGTCTG**ttgaatagtggggcataactca**

### ***LUMINIDEPENDENS*** **(*LD*) from *Zea mays*, 470 bp final amplicon**
(size distinguishable from V4 or V3V4 16S rDNA primers)

*Underlined letters represent the universal overhang.*
*Bracketed grey boxed letters represent "linkers" - sequences designed not to anneal to the template and reduce unintended binding of the rest of the primer.*
*Lowercase letters are template-specific.*
*"F1", "bcGA", "bcTC", "bcAG",* and *"bcCT" forward primers perform identically but each introduces different sequenced bases which serve as an additional barcode. See (Lundberg et al., 2013).*

**FORWARD:**
>Zm_LD_bcGA_G-47184
TCCCTACACGACGCTCTTCCGATCT GA **[TA]** **tctgccctggtggagtacccat**
>Zm_LD_bcTC_G-47185
TCCCTACACGACGCTCTTCCGATCT TC **[TA]** **tctgccctggtggagtacccat**
>Zm_LD_bcAG_G-47186
TCCCTACACGACGCTCTTCCGATCT AG **[TA]** **tctgccctggtggagtacccat**
>Zm_LD_bcCT_G-47187
TCCCTACACGACGCTCTTCCGATCT CT **[TA]** **tctgccctggtggagtacccat**

**REVERSE:**
>Zm.LD_R1_G-47158
GGAGTTCAGACGTGTGCTCTTCCGATCT **cctcggcgcctccgggtt**

>Z.mays_LD_sequenced_region_375bp_final_amplicon_512bp
**gccctggtggagtacccat**GGGACGGGCACCACCAGCGGCACTCGAGGTCACCAGATCCTGGCGTGGTCCGGG
ACTACGACACCGACTATGGCGGTGCGCAGGGCTACAGTCAGCAGCCCCTGACGCAGTGGAGTGCAGGGAAAGT
GCAGCAGCAGGGCTACAATCCTGAGCCGTCAAGGCAGTGGAGTTCTTCCCAGGCGCACCAGGGCGGCTACGCA
CCCGCCGAGCCGTCGAGGCAGTGGAGTTCTTCCCAGGCGCACCAGAGCTACGCTCCCGAGCTACCGAGGCAGT
GGAGCTCCGAACGCCGTGGCTACGATGATGCGGAGCCCTCGAGGACATGGAGCTCCGGCCAGCAG**aacccgga
ggcgccgagg**

### Note on Adapting Tagging Sets for Other Protocols

For hamPCR to work efficiently, the tagging primers should not have a tendency to dimerize. Simply changing the universal overhangs may be sufficient to use hamPCR with other two-step PCR protocols. *The following are starting suggestions only, and have not been tested experimentally.*

To adapt a forward tagging primer to that in (Gohl et al., 2016), replace the **upper-case, unbracketed letters** in the forward primer for that template (shown in orange in the example below) with "TCGTCGGCAGCGTCAGATGTGTATAAGAGACAG".

For example, for "**At.GI_F1_G-46602**" for *A. thaliana GI*,
"TCCCTACACGACGCTCTTCCGAT**ct** **gtaaagataaatgggtcatctaa**" would become:
"TCGTCGGCAGCGTCAGATGTGTATAAGAGACAG **ctgtaaagataaatgggtcatctaa**"

To adapt a reverse tagging primer to that in (Gohl et al., 2016), replace the **upper-case, unbracketed letters** (shown in orange in the example below) in the reverse primer for that template with: "GTCTCGTGGGCTCGGAGATGTGTATAAGAGACAG".

For example, for "**At.GI_R502bp_G-46614**" for *A. thaliana GI*,
"GGAGTTCAGACGTGTGCTCTTCCGATCT **tccttctgaaccggtgtattc**" (this work) would become:
"GTCTCGTGGGCTCGGAGATGTGTATAAGAGACAG **tccttctgaaccggtgtattc**".

Rather than adapt the forward tagging primers to work with the forward PCR primer in (Lundberg et al., 2013), we recommend to use the forward primer here. The reverse tagging primers can be used directly with the reverse PCR primers in (Lundberg et al., 2013).

### Primers: Exponential PCR

For the full PCR primer set, please see Supplementary Table 1. PCR primers should be HPLC-purified or purified with an alternative method to reduce truncated primers.

**FORWARD**

```
>PCR_F_G-40610
AATGATACGGCGACCACCGAGATCTACACTCTTTCCCTACACGACGCTCTTC
```

**REVERSE:**

Example PCR_R_indexed primers. Same set as that published in (Lundberg et al., 2013).

| **Model Reverse Primer** | **CAAGCAGAAGACGGCATACGAGAT XXXXXXXXX GTGACTGGAGTTCAGACGTGTGCTC** |
|---|---|
| PCR R bc1 | CAAGCAGAAGACGGCATACGAGAT TTACCGACG GTGACTGGAGTTCAGACGTGTGCTC |
| PCR R bc2 | CAAGCAGAAGACGGCATACGAGAT ATTGGACAC GTGACTGGAGTTCAGACGTGTGCTC |
| PCR R bc3 | CAAGCAGAAGACGGCATACGAGAT TCGCATGGA GTGACTGGAGTTCAGACGTGTGCTC |
| Etc. | |

### Supplementary Figure 2

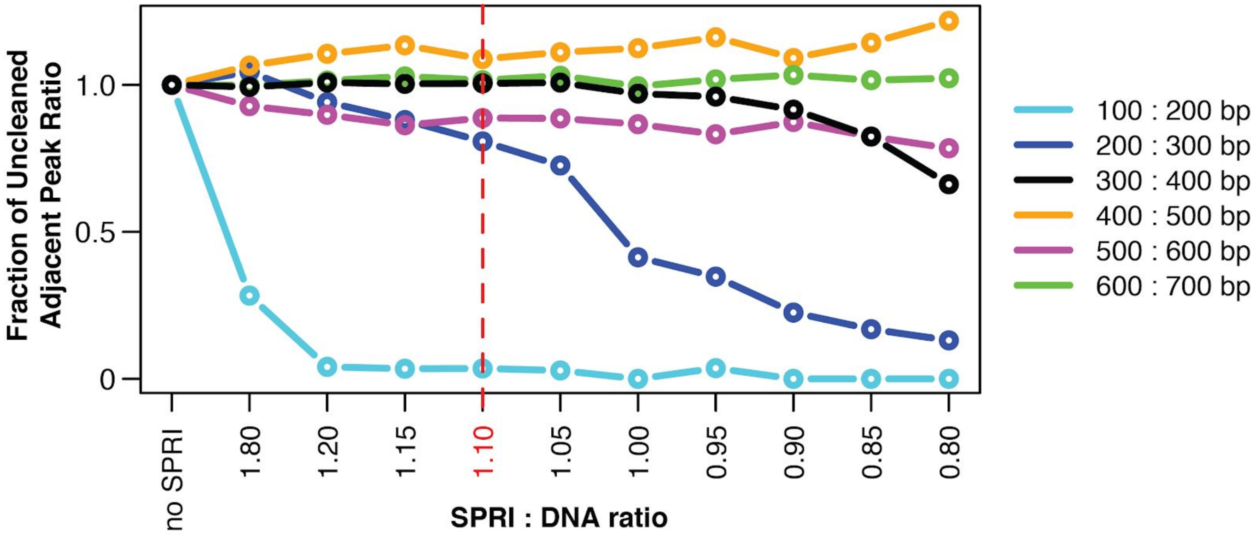


**Supplementary Figure 2. SPRI ratios |** SPRI beads in a polyethylene glycol (PEG) solution, such as AMPure XP, preferentially bind longer DNA fragments, making them useful for removing free primers and primer dimers from reactions. As the PEG concentration decreases, a wider range of short fragments can no longer bind to the beads, and primer dimers are more completely eliminated. However, if the PEG concentration is too low, the range of fragment sizes eliminated could include DNA of interest. For hamPCR, it is important that the SPRI cleanup does not affect the ratio of the host and microbial amplicons, which could lead to systematic bias and noise. To determine an acceptable PEG concentration, we tested cleanups with SPRI (AMPure XP) beads at different SPRI : DNA ratios (resulting in a range of PEG concentrations) on a standard DNA ladder (GeneRuler DNA Ladder Mix, Thermo Scientific, Waltham, MA, USA), and quantified abundance of the purified fragments with a Bioanalyzer (Agilent, Santa Clara, CA, USA). Using the pure, uncleaned ladder, we calculated the ratio of each peak's abundance to the adjacent larger peak (200 : 300 bp, 300 : 400 bp, etc.), and used this set of abundance ratios as a baseline (no SPRI, far left). With each successive decrease in SPRI : DNA ratio (from left to right), we looked for a decrease in the abundance ratio between adjacent bands, which would indicate elimination of the smaller fragment. Of highest interest is the 300 : 400 bp ratio (black), because the smallest tagged templates in hamPCR are around 300 bp. We determined SPRI : DNA ratios less than 1.0 endangered the 300 : 400 ratio, and thus decided to conservatively use a SPRI : DNA solution of 1.1 : 1 or higher for the cleanup of tagged products.

### Supplementary Figure 3

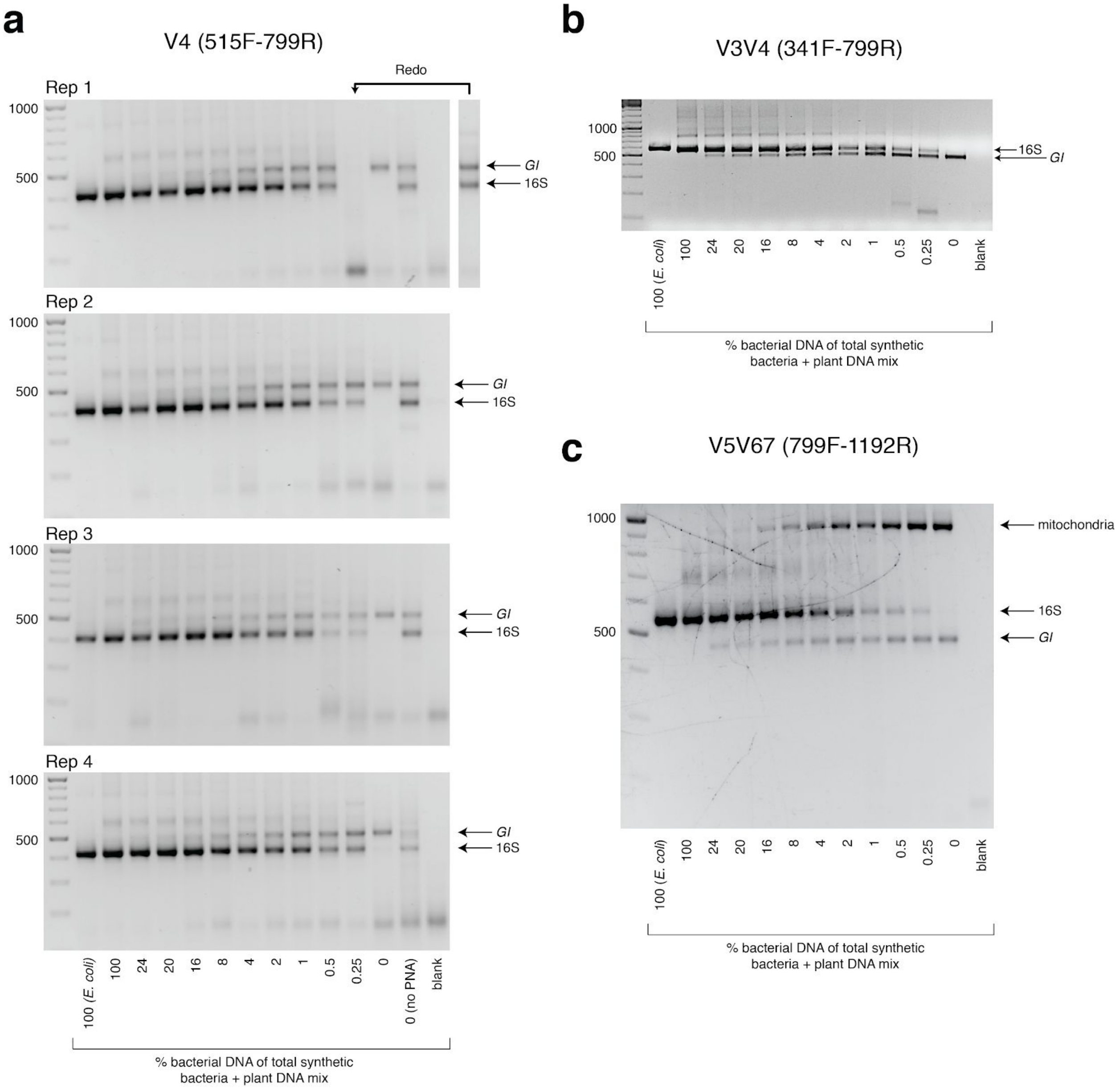


**Supplementary Figure 3. Gel images from synthetic titration panel | a**, hamPCR using the 502 bp *A. thaliana GI* amplicon and the ~420 bp V4 16S rDNA amplicon (515F - 799R) was performed with the samples of the the synthetic titration panel, and 5 µL of the PCR product from each sample was analyzed on a 2% agarose gel. This was done across four replicates each prepared with an independent PCR master mix. Due to a technical failure in replicate 1, the sample with 0.25% bacteria did not work and was redone in a separate reaction. **b**, The same as (a), except using the ~587 bp V3V4 16S rDNA amplicon (341F - 799R). **c**, The same as in (a) and (b), except using the 466 bp *A. thaliana GI* amplicon and the ~540 bp V5V6V7 16S rDNA amplicon (799F - 1192R). This 16S rDNA primer set also amplifies a ~770 bp region of mitochondria, producing a final amplicon of about 900 bp. In practice, the mitochondrial band is usually removed by gel extraction of the desired amplicons prior to sequencing.

### Supplementary Figure 4

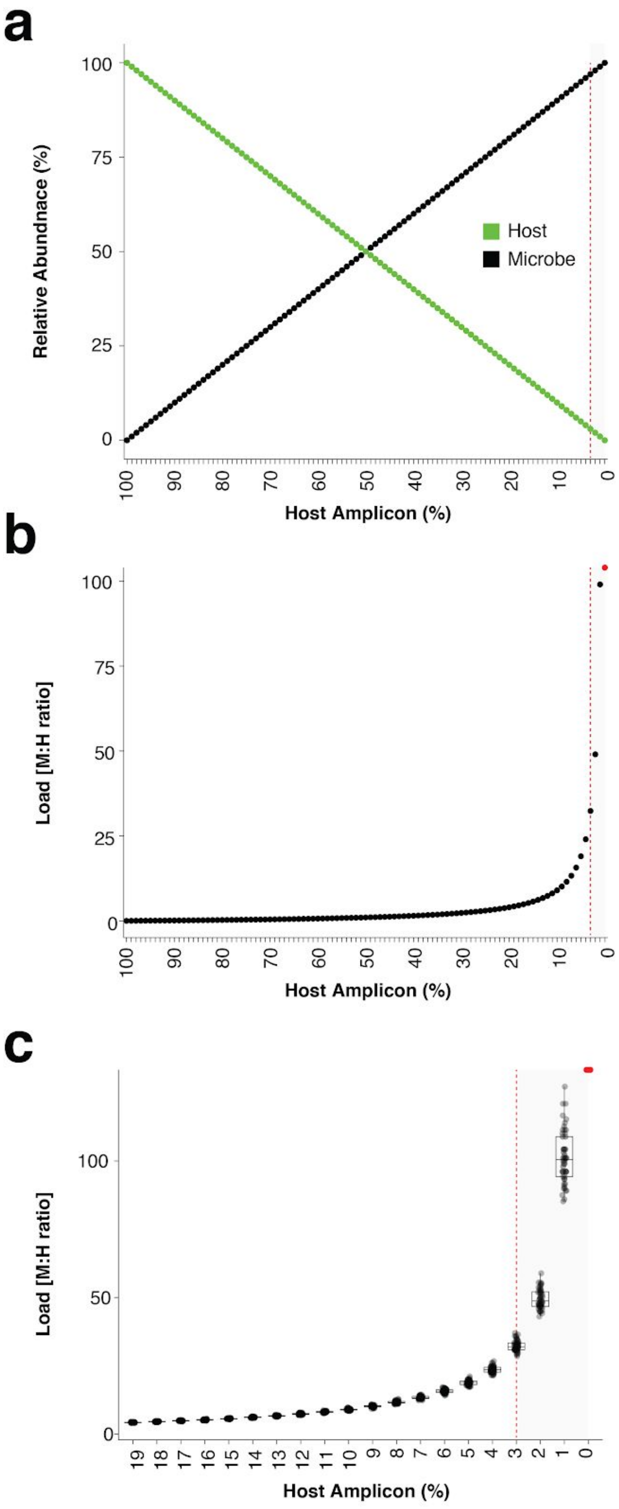


**Supplementary Figure 4. In-silico load simulations | a**, One hundred and one samples were simulated by combining 0 to 100 microbial sequence counts (black points) with 100 to 0 sequence counts (green points). The x-axis shows the percentage of the sample occupied by host amplicon, decreasing from left to right. The vertical red dotted line indicates the position of 3% host amplicon abundance. **b**, The microbial sequence counts from (a) were converted to microbial load by dividing by the host sequence counts. Note that as the host amplicon abundance approaches 0, the microbial load climbs towards infinity. The red point at 0% host amplicon abundance represents infinite load. The vertical red dotted line indicates 3% host amplicon abundance. **c**, The 100 plant and microbial sequence counts for each sample in (a) were multiplied by 10,000 to make virtual samples with 1 million sequence counts. These were then subsampled 50 times each to 10,000 reads to simulate random sampling noise. Microbial load was calculated as in (b) by dividing microbial counts by host counts using the 10,000 count subsampled data. Only host amplicon percentages from 19% to 0% are shown to zoom in on low host abundances. The red point at 0% host amplicon abundance represents infinite load. The vertical red dotted line indicates the position of 3% host amplicon abundance. Note that not only does load climb quickly towards infinity as host counts approach 0, but also sampling noise has a greater impact on microbial load, leading to greater variance.

### Supplementary Figure 5

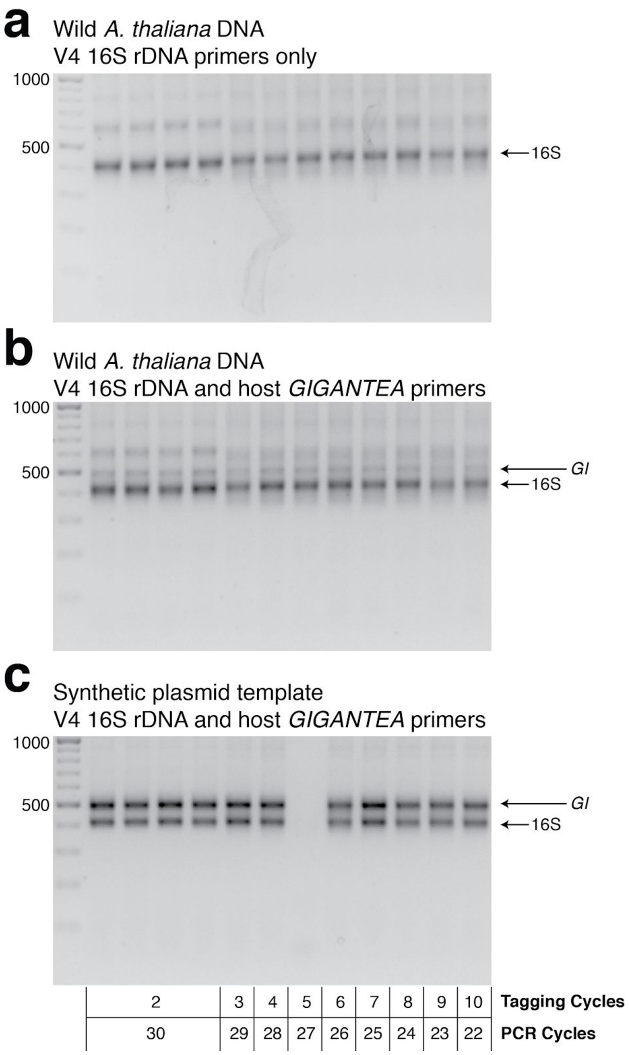


**Supplementary Figure 5. Gel of template tagging cycling tests | a**, hamPCR using *only* primers for the V4 region of the 16S rDNA (primers 515F - 799R) was run on pooled DNA made from wild *A. thaliana* leaves. This represents the output of common 2-step 16S rDNA amplification protocols. Four replicates were produced with 2 cycles of the tagging reaction and 30 cycles PCR, and additional replicates were produced with 3-10 tagging cycles paired with 29 to 22 PCR cycles (for a total of 32 cycles), as shown under panel C. **b**, The same DNA, PCR master mix, and cycling conditions as in (a), except with the addition of primers for the 502 bp *A. thaliana GI* amplicon. **c**, The same as in (b), except with a synthetic plasmid template containing a fragment of the *A. thaliana GI* gene and 16S rDNA from *Pto* DC3000 in a 1:1 ratio. Tagging cycles and PCR cycles for each lane are shown beneath.

### Supplementary Figure 6

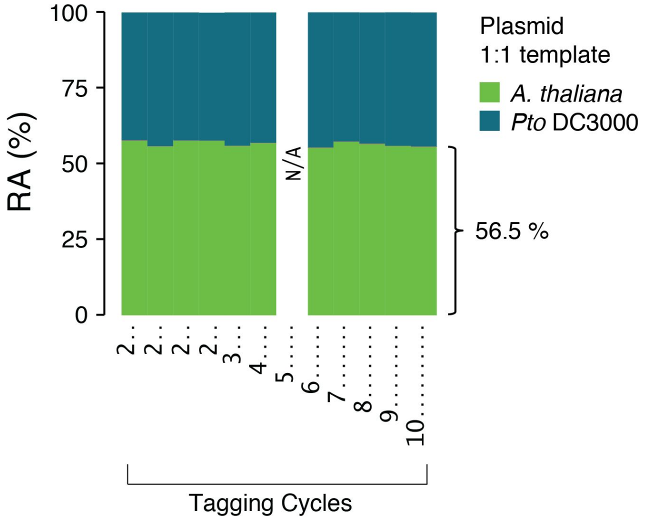


**Supplementary Figure 6. Effect of varying tagging cycles on the plasmid 1:1 host:microbe template |** A synthetic plasmid-borne template including the *Pto* DC3000 16S rDNA and a portion of the *A. thaliana GI* gene was used as template for hamPCR using the V4 region of the 16S rDNA (primers 515F - 799R) and the 502 bp *A. thaliana GI* amplicon. Four replicates were produced with 2 cycles of the tagging reaction and 30 cycles PCR, and additional replicates were produced with 3-10 tagging cycles paired with 29 to 22 PCR cycles (for a total of 32 cycles). A technical error produced no data for 5 tagging cycles (see Supplementary Figure 4). The number of tagging cycles did not influence the ratio of host to microbe, and the host amplicon was slightly, but consistently overrepresented at 56.5% of the total sequence in these samples.

### Supplementary Figure 7

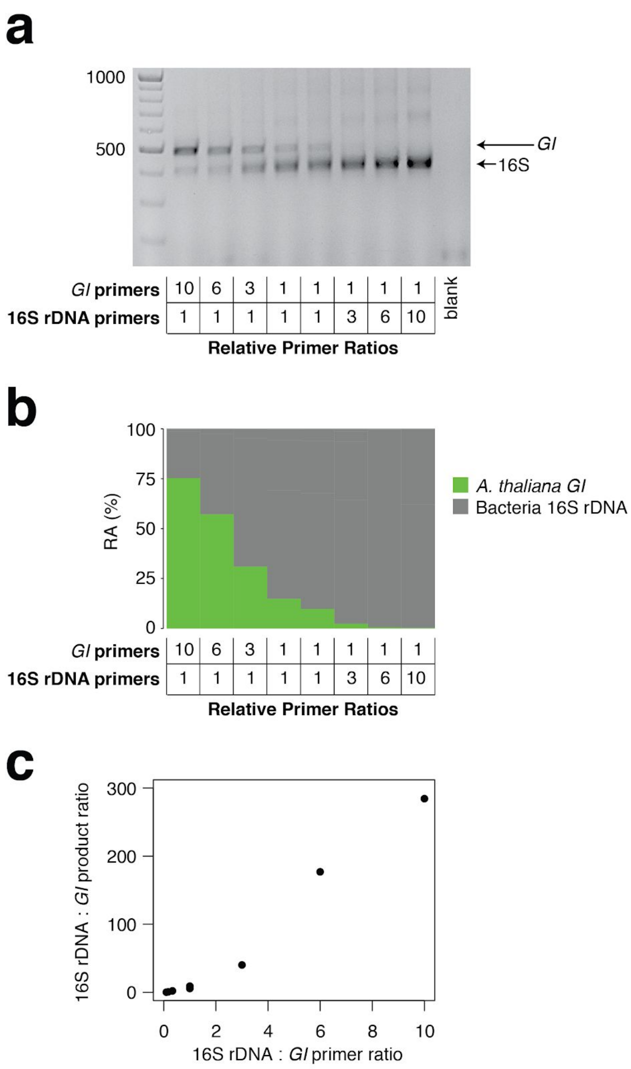


**Supplementary Figure 7. Effect of primer pair concentration on product |** The same wild *A. thaliana* DNA pool was amplified with hamPCR using 8 different ratios of the 16S rDNA primer pair to the *GI* primer pair, ranging from 1/10 to 10. **a**, 2% agarose gel of the prepared libraries **b**, Relative abundance (RA) of bacteria and host amplicons in the sequence data. **c**, Scatterplot of microbe to host product ratios plotted against the microbe to host primer ratios used to produce them. A ten-fold difference in primer ratios resulted in an over 200-fold difference in product ratios, underscoring the importance of steady primer ratios (through the use of a mastermix of primers) across an experiment.

### Supplementary Figure 8

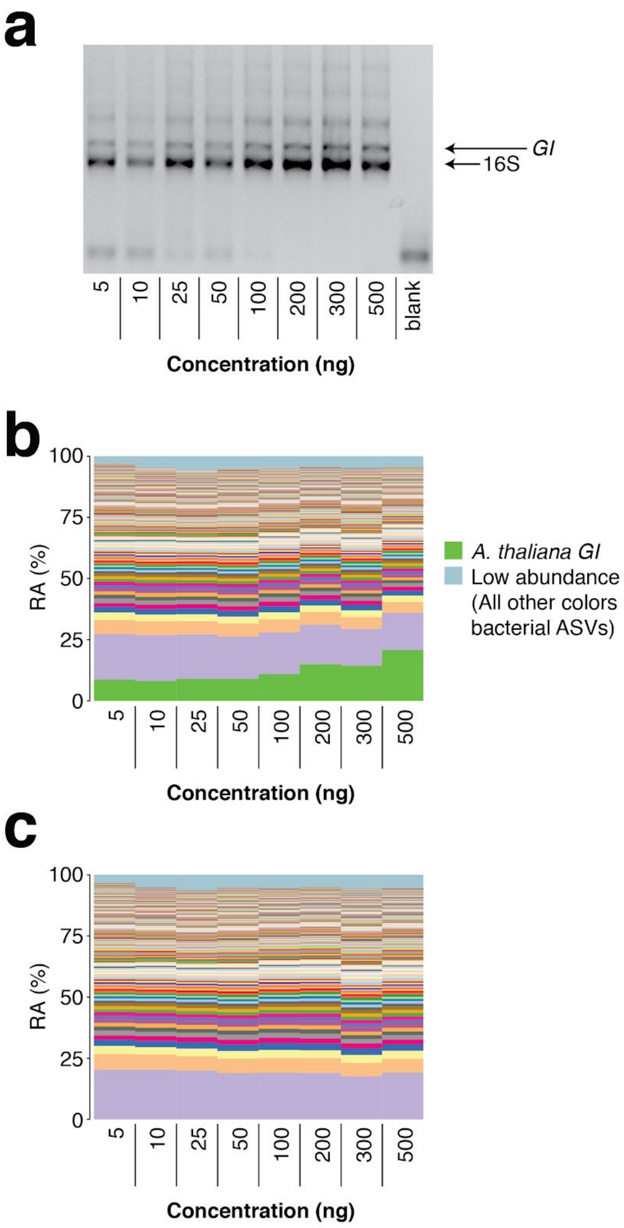


**Supplementary Figure 8. Total template concentration test |** A panel of 8 concentrations of wild *A. thaliana* leaf DNA, ranging from 5 to 500 ng per reaction (approximately ($3.6\times10^4$ to $3.6\times10^6$ host *GI* template copies per reaction assuming a 135 Mb *A. thaliana* genome), were made into hamPCR libraries. **a**, 2% agarose gel of the prepared libraries **b**, Relative abundance (RA) of all amplicons in the reaction. The *A. thaliana GI* ASV appears to increase at the highest template concentrations, but remains of constant abundance through the standard template range of 5 to 100 ng. **c**, The *A. thaliana GI* ASV has been removed and the bacterial ASVs have been rescaled to give bacterial relative abundance.

### Supplementary Figure 9

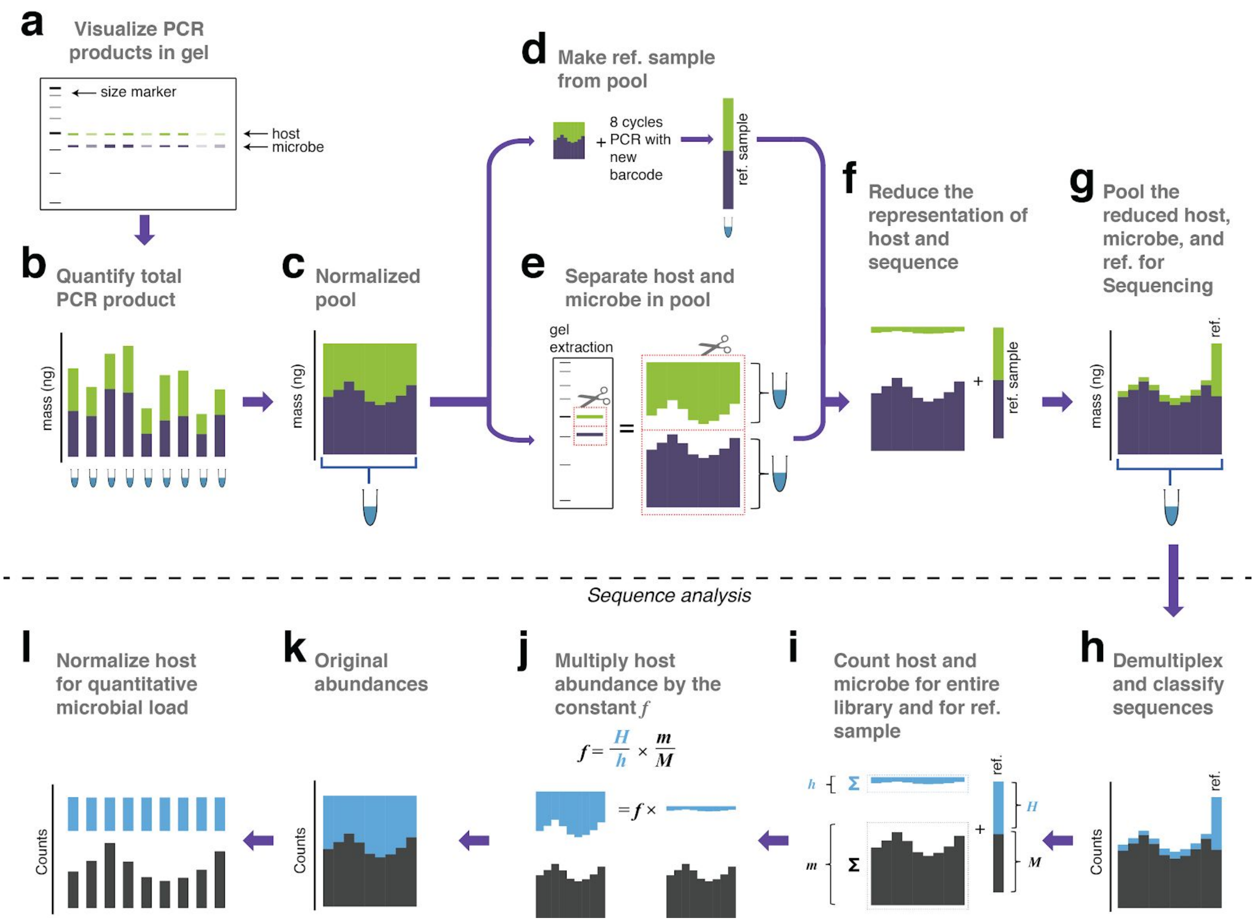


**Supplementary Figure 9. Library remixing prior to sequencing | a**, PCR products analyzed on a 2% agarose gel. Because the host and microbe amplicons are different lengths, they migrate at different speeds and can be independently resolved. **b**, The total DNA concentration of each PCR product is measured. **c**, The samples are pooled at equimolar ratios into a single tube. **d**, A very small aliquot of the pool from (c) is re-amplified with 8 cycles of PCR to replace all barcodes in the pool with a new barcode, creating a reference sample which is set aside. **e**, A large aliquot of the pool from (c) is run on an agarose gel to physically separate the host and microbial fractions. These fractions are separately purified and quantified. Because the extraction is done on the pool, samples are not differentially affected by extraction inefficiencies. **f**, The DNA concentrations in the host fraction, microbial fraction, and reference sample are each quantified, and the desired molarity of each is aliquoted. Here, host amplicons are depleted to approximately 5% of the total library. The actual mixing ratio is not important, because the reference sample preserves the correct ratio of host to microbe. **g**, The reduced host fraction, microbial fraction, and the reference sample are pooled for sequencing. **h**, All sequences are quality filtered, demultiplexed, and taxonomically classified using the same parameters. **i**, Host and microbe sequence counts are summed from the samples comprising the pooled library (*h* and *m* respectively), and from the reference sample (*H and M respectively*). **j**, *H, h, M, and m* are used to calculate the scaling constant *f* for the dataset. All host sequence counts are multiplied by *f* to reconstruct the original microbe to host ratios. **k**, Reconstructed original abundances (compare to original abundances in B. **l**, Finally, each sample is divided by host abundance to normalize host counts, resulting in microbial sequence counts that represent the microbial load of each micobe on the host.

### Supplementary Figure 10

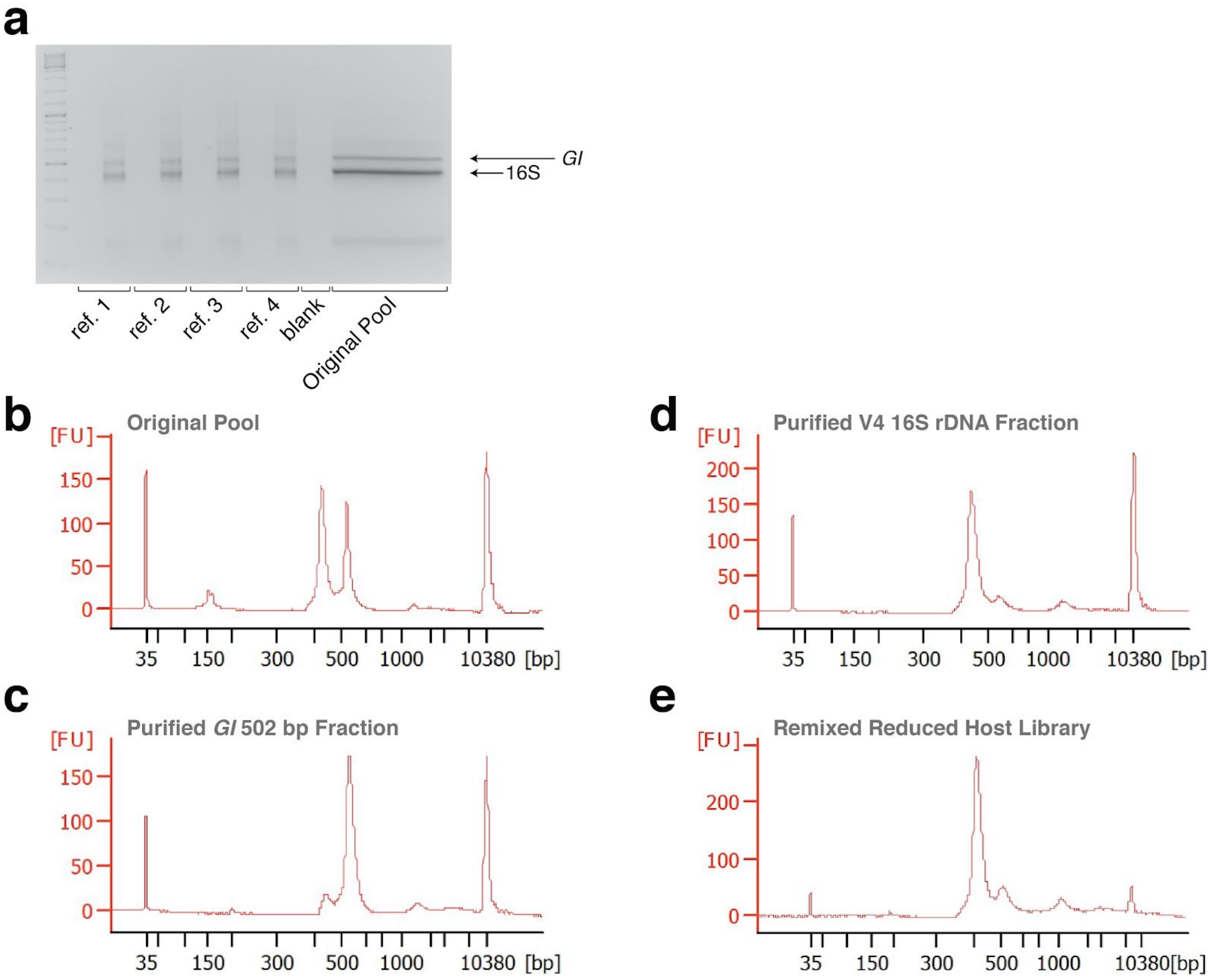


**Supplementary Figure 10. Gels and Bioanalyzer traces showing steps of remixing | a**, Gel showing four replicate reference samples made from the original pool, confirming successful re-barcoding. For each pair of lanes, the left lane is 5 µL of the PCR reaction prior to 8 cycles of amplification, and the right lane is 5 µL after cycling. Size separation of the original pool for purification of the host (*GI*) and microbial (16S) fractions is shown in the wide lane. **b**, Bioanalyzer trace of the original pool, showing nearly equally abundant host and microbial fractions. **c**, Bioanalyzer trace of the purified host fraction. Some host fraction has been inadvertently and harmlessly co-purified. **d**, Bioanalyzer trace of the purified host fraction. A small amount of the microbial fraction has been inadvertently and harmlessly co-purified. **e**, Bioanalyzer trace of the remixed reduced host library.

### Supplementary Figure 11

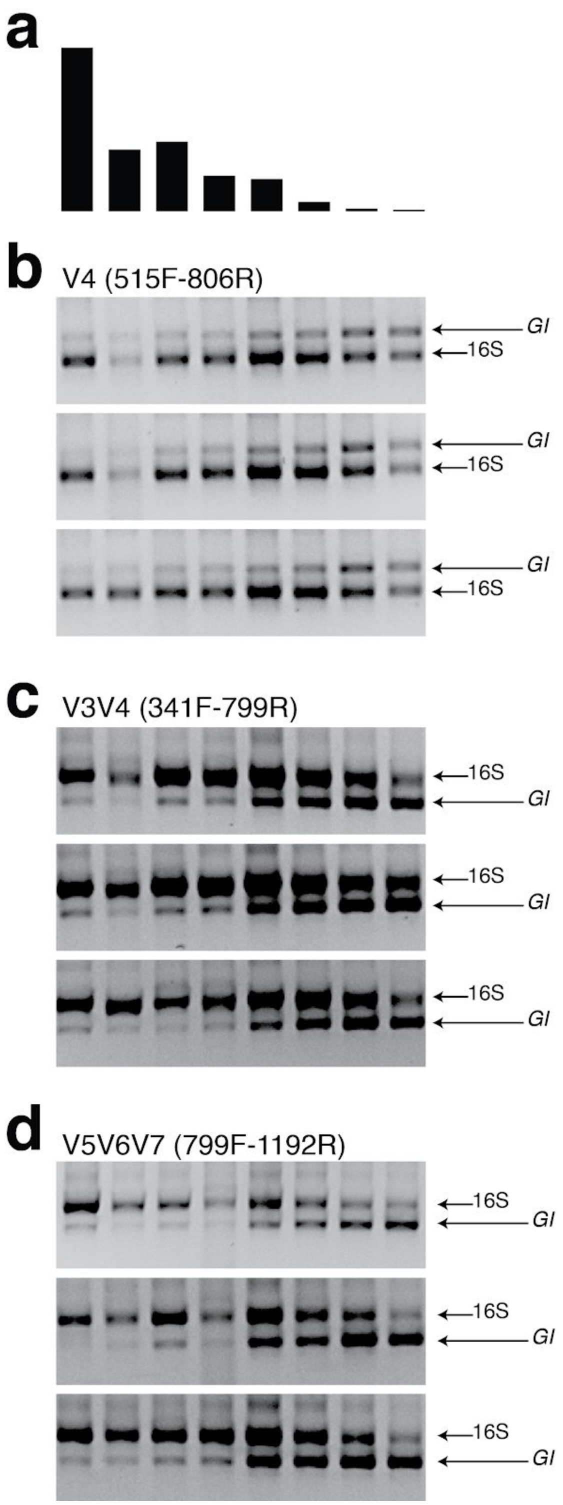


**Supplementary Figure 11. Gel pictures from hamPCR applied to wild *A. thaliana* leaf DNA samples | a**, The bacterial load from 8 wild *A. thaliana* samples (Regalado et al., 2020), as determined by shotgun sequencing. **b**, Three replicates of hamPCR products made from the same 8 DNA samples, using the 502 bp *A. thaliana* *GI* amplicon and the ~420 bp V4 16S rDNA amplicon (515F - 806R). The 806R primer was used here instead of the 799R primer used elsewhere in the manuscript so that the 16S rDNA data could be directly compared to previous V4 16S rDNA amplicon data from those samples, which also used 515F - 806R. The products were visualized on a 2% agarose gel. **c**, Same as (b), but hamPCR was performed using the ~590 bp V3V4 16S rDNA amplicon (341F - 799R). **d**, Same as in (b) and (c), but hamPCR was performed using the 466 bp *A. thaliana* *GI* amplicon and the ~540 bp V5V6V7 16S rDNA amplicon (799F - 1192R). Note that in (c) and (d), the bacterial amplicon is longer than the host amplicon.

### Supplementary Figure 12

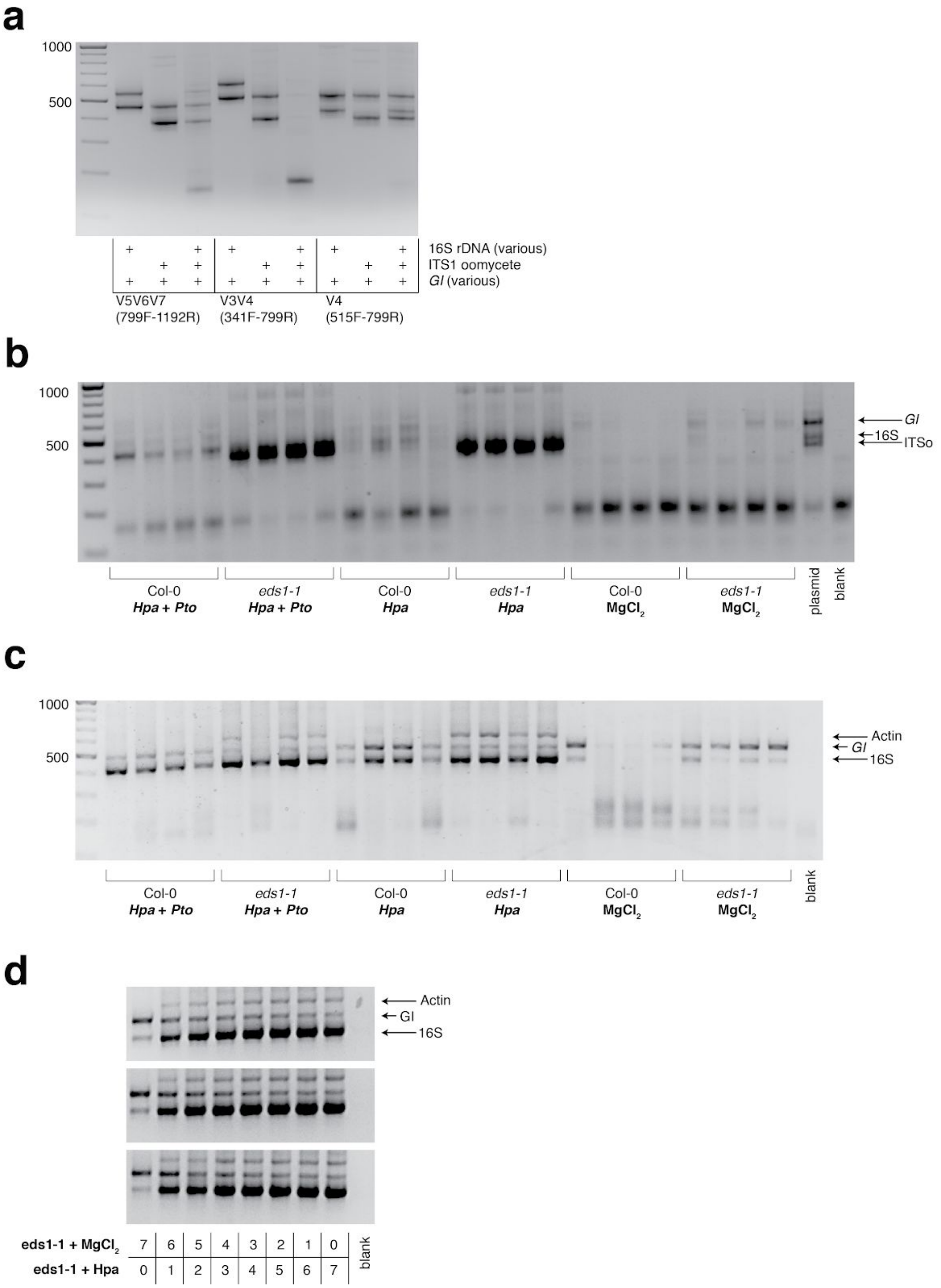

**Supplementary Figure 12. Gel pictures of hamPCR applied to one host and two microbial amplicons | a**, Test of primer compatibility for primer pairs targeting plant and bacteria, plant and oomycete, or plant and bacteria and oomycete, for the ITSo rDNA amplicon, three alternate bacterial amplicons, and two plant amplicons as shown. Some combinations perform better than others and produce fewer dimers. Shown on a 2% agarose gel. **b**, The samples of the *Hpa* and *Pto* DC3000 co-infection using ITSo rDNA primers, V4 16S rDNA primers, and 502 bp *A. thaliana GI* primers. The signal from the ITSo primers overwhelmed the reaction. **c**, hamPCR library of the *Hpa* and *Pto* DC3000 co-infection as described in Figure 5a-c using *Hpa* actin primers, V4 16S rDNA primers, and 502 bp *A. thaliana GI* primers. **d**, hamPCR libraries made with the titration of plant DNA infected with *Hpa* as described in Figure 5d, using actin primers, V4 16S rDNA primers, and 502 bp *A. thaliana GI* primers.

### Supplementary Figure 13

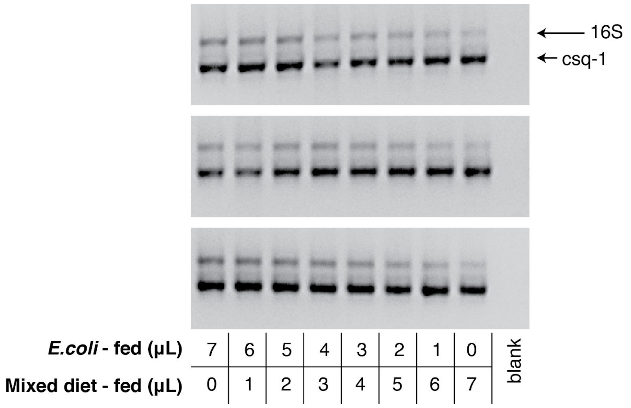


**Supplementary Figure 13. Gel of *P. pacificus* hamPCR titration libraries |** DNA was prepared from two populations of *P. pacificus* strain PS312, one fed with only *E. coli* and another fed with *E. coli*, *P. syringae*, and *Lysinibacillus xylanilyticus* (mixed diet). The two DNA samples were each adjusted to 6 ng/μL and titrated into each other using the ratios shown below the gel. The 2% agarose gel shows resulting hamPCR libraries using the ~540 bp V5V6V7 16S rDNA amplicon (primers 799F - 1192R) and the 470 bp *P. pacificus calsequestrin* (*csq-1*) amplicon.

### Supplementary Figure 14

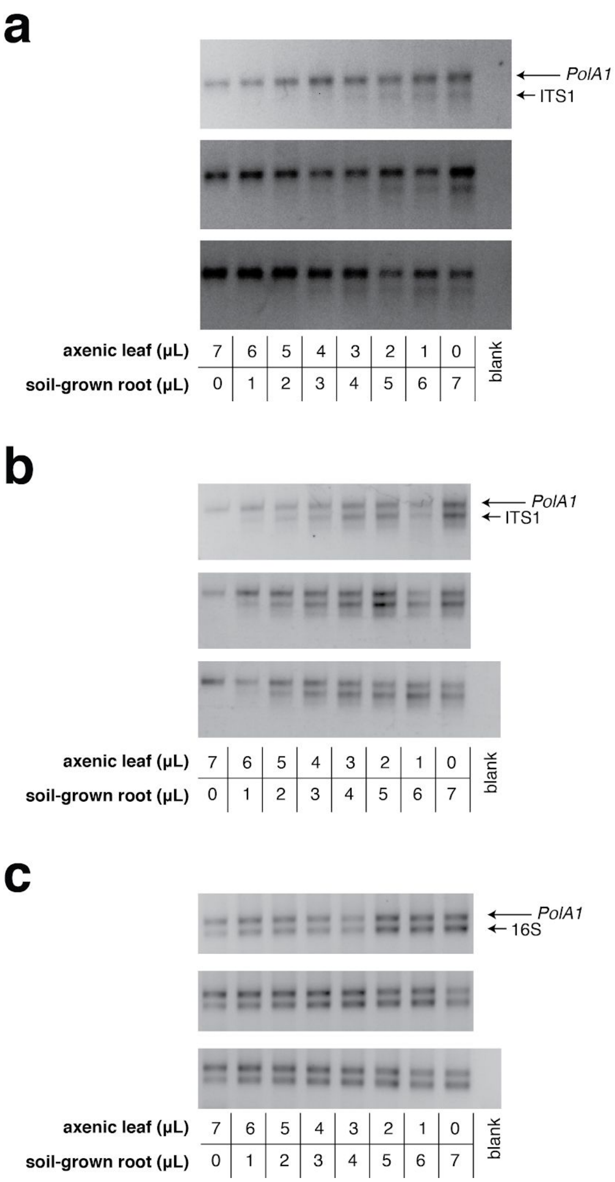


**Supplementary Figure 14. Gels of *T. aestivum* hamPCR titration libraries | a-c:** DNA was made from axenic *T. aestivum* leaf DNA and surface-sterilized *T. aestivum* roots that had been cultivated outdoors in potting soil. The two DNA samples were each adjusted to ~ 60 ng/µL and titrated into each other using the ratios shown below the gel. hamPCR products are displayed on a 2% agarose gel. **a**, hamPCR products using the ITS1 fungi amplicon and the 497 bp *T. aestivum PolA1* amplicon. The ITS1 and *PolA1* primer pairs were mixed in a 1:1 ratio. Tagging was performed with 2 cycles and PCR was performed with 30 cycles. Note that ITS1 amplicons form a smear on the gel due to a wide range of amplicon sizes. **b**, Same as (a), but the ITS1 and *PolA1* primer pairs were mixed in a 2:1 ratio to favor ITS1 amplification. To help amplification of lower quantities of tagged templates, tagging was performed with 7 cycles and PCR was performed with 25 cycles. **c**, hamPCR products using primers for V4 16S rDNA from bacteria (primers 515F - 799R) and the 497 bp *T. aestivum PolA1* amplicon. A standard protocol of 2 tagging cycles and 30 PCR cycles was used.

### Supplementary Figure 15

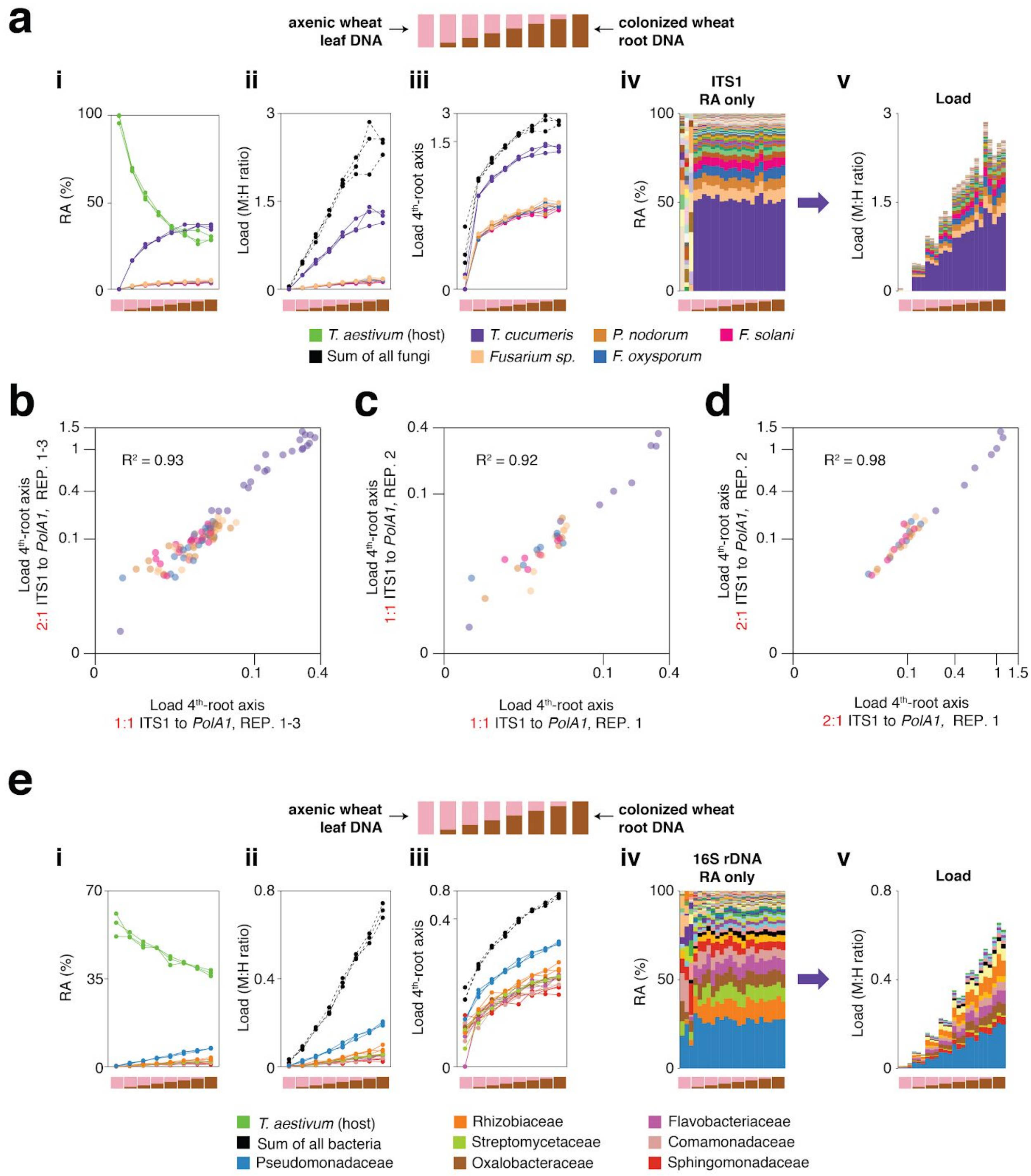


**Supplementary Figure 15. ITS1 load in *T. aestivum* with a 2:1 microbe to host tagging primer ratio, and 16S rDNA load in *T. aestivum* | a**, Similar to Figure 5f in the main text, axenic *T. aestivum* leaf DNA was titrated into DNA from *T. aestivum* roots that had been cultivated outdoors in potting soil to make a panel of eight samples simulating different levels of infection. hamPCR was performed for three replicates per sample using ITS1 fungal primers and *T. aestivum PolA1* primers. Unlike Figure 5f, the ITS1 and *PolA1* primer pairs were mixed in a 2:1 ratio to favor ITS1 amplification, and tagging was performed with 7 cycles and PCR was performed with 25 cycles. **i:** the relative abundance of hamPCR amplicons with median abundance above 0.15%. **ii:** after using host ASV to convert amplicons to load. The cumulative load is shown in black. **iii:** the load on a 4th-root transformed y-axis, showing less-abundant families. **iv:** stacked column visualization of all ASVs for the panel as it would be seen with pure ITS1 data. **v:** stacked-column plot of the panel corrected for microbial load. **b**, Loads for abundant fungal ASVs following hamPCR with a 2:1 ITS1 to *PolA1* primer ratio plotted against loads for the same ASVs following hamPCR with a 1:1 ITS1 to *PolA1* primer ratio (as displayed in Figure 5f). Shown on 4th-root axes. $R^2$, coefficient of determination. The correlation comparing ASVs in 1:1 and 2:1 ITS1 to *PolA1* primer ratios is just as high as for the correlation between technical replicates with the same priming conditions, shown next in (c) and (d). **c**, Same as (b), but with one technical replicate of hamPCR with a 1:1 ITS1 to *PolA1* primer ratio plotted against a second technical replicate of the 1:1 ratio. **d**, Same as (b) and (c), but for one technical rep of hamPCR with a 2:1 ITS1 to *PolA1* primer ratio plotted against a second technical replicate of the 2:1 ratio. **e**, hamPCR using the V4 16S rDNA bacterial amplicon and the *T. aestivum PolA1* amplicon, using a standard 1:1 microbe to host primer ratio and normal cycling. Subpanels i-v as in (a), with bacterial families shown instead of fungal ASVs.

### Supplementary Figure 16

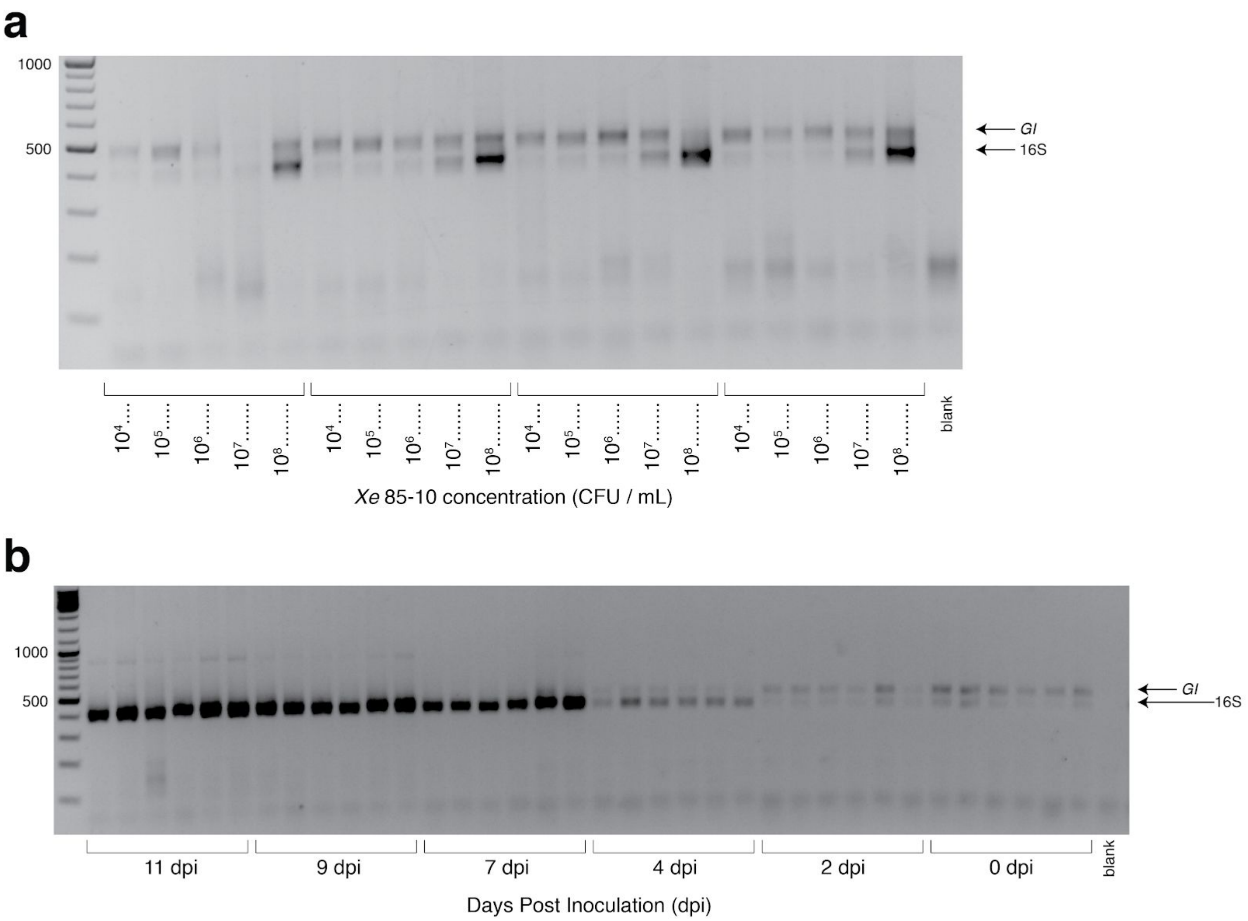


**Supplementary Figure 16. Gels from *C. annuum* experiments | a)** 2% agarose gel showing hamPCR libraries for 4 replicates of *C. annuum* infiltration with *Xanthomonas euvesicatoria* (*Xe*) 85-10. The tagging step was done with V4 16S rDNA primers (515F - 799R) and *C. annuum GI* primers. **b)** 2% agarose gel showing hamPCR libraries for 6 replicates per timepoint of a *C. annuum* growth curve following infiltration with $10^4$ CFU/mL *Xe* 85-10.

### Supplementary Figure 17

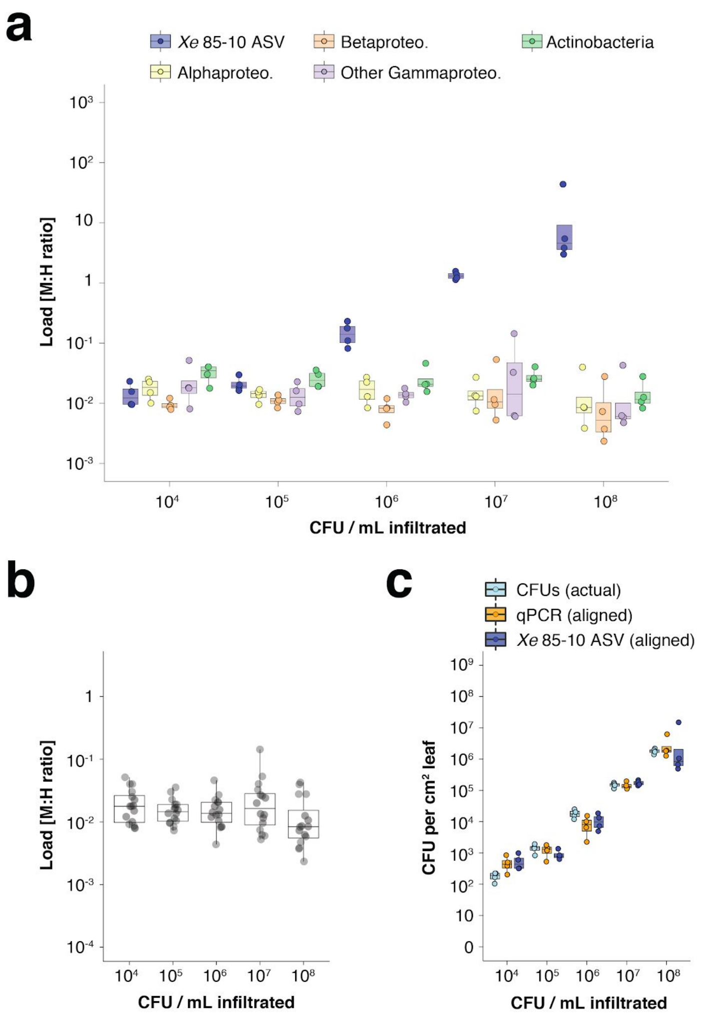


**Supplementary Figure 17. hamPCR can be generalized to *Xanthomonas* infection of *C. annuum* | a-c**, Infiltration experiment. All y-axes are on a base-10 logarithmic scale. In all boxplots, the median is represented by a horizontal line and box boundaries show the lower and upper quartiles. Whiskers extend from the box up to 1.5 times the interquartile range. **a**, *Xe* 85-10 was inoculated into *C. annuum* leaves at five concentrations, leaves were harvested immediately afterwards, and hamPCR was performed. The bacterial load is shown for the particular ASV corresponding to *Xe* 85-10, as well as for the major bacterial classes. **b**, The total load for all bacterial classes shown in (a) at each concentration. **c**, Actual CFU counts for *Xe* 85-10 in the infiltration experiment juxtaposed with aligned qPCR and hamPCR loads.

### Supplementary Discussion: Synthetic Template

The ITS1 region from *Agaricus bisporus* (3) was amplified with the forward (1) and reverse (2) primers below:

>1___ITS1_forward_for_ITS_G-46643
AGAAGGAGATATACCATGG**cttggtcatttagaggaagtaa**
>2___ITS-GI_reverse_for_ITS_G-46644
ACAGGTTGATTCGCCTCAATA**gctgcgttcttcatcgatgc**
>3___ITS1_Agaricus_bisporus
**cttggtcatttagaggaagtaa**AAGTCGTAACAAGGTTTCCGTAGGTGAACCTGCGGAAGGATCATTATTGAATTATGTTTTCTAGATGGGTTGT
AGCTGGCTCTTCGGAGTATGTGCACGCCTGTCTGGACTTCATTTTCATCCACCTGTGCACCTTTTGTAGTCTTTTTCAGGTATTGGAGGAAGTG
GTCAGCCTATCAGCTCTTTGCTGGATGTAAGGACTTGCAGTGTGAAAACAGTGCTGTCCTTTACCTTGGCCATGGAATCTTTTTCCTGTTAGAG
TCTATGTTATTCATTATACTCTTAGAATGTCATTGAATGTCTTTACATGGGCTATGCCTATGAAAATTATTATACAACTTTCAGCAACGGATCT
CTTGGCTCTC**gcatcgatgaagaacgcagc**

A portion of the *GIGANTEA* (*GI*) gene from *Arabidopsis thaliana* (3) was amplified with the forward (1) and reverse (2) primers below:

>1___ITS-GI forward for_GI_gene_G-46645
GCATCGATGAAGAACGCAGC**tattgaggcgaatcaacctgt**
>2___GI-16S_reverse_for_GI_gene_G-46646
CTGAGCCAGGATCAAACTCT**cctgagccttcatctgaatgt**
>Gigantea_partial_Arabidopsis
**tattgaggcgaatcaacctgt**ATCTAACAATCAAACTGCTAACCGTAAAAGTAGGAATGTCAAGGGACAGGGACCTGTGGCAGCATTTGATTCAT
ACGTTCTTGCTGCTGTTTGTGCTCTTGCCTGTGAGGTTCAGCTGTATCCTATGATCTCTGGTCGGGGAACTTTTCCAATTCTGCCGTGGCTGG
AACTATTACAAAGCCTGTAAAGATAAATGGGTCATCTAAAGAGTATGGAGCTGGGATTGACTCGGCAATTAGTCATACGCGCCGAATTTTGGCA
ATCCTAGAGGCACTCTTTTCATTAAAACCATCTTCTGTGGGGACTCCATGGAGTTACAGTTCTAGTGAGATAGTTGCTGCGGCCATGGTTGCAG
CTCATATTTCCGAACTGTTCAGACGTTCAAAGGCCTTGACGCATGCATTGTCTGGGTTGATGAGATGTAAGTGGGATAAGGAAATTCATAAAAG
AGCATCATCATTATATAACCTCATAGATGTTCACAGCAAAGTTGTTGCCTCCATTGTTGACAAAGCTGAACCCTTGGAAGCCTACCTTAAGAAT
ACACCGGTTCAGAAGGATTCTGTGACCTGTTTAAACTGGAAACAAGAGAACACATGTGCAAGCACCACATGCTTTGATACAGCGGTGACATCCG
CCTCAAGGACTGAAATGAATCCAAGAGGAAACCATAAGTATGCTAG**acattcagatgaaggctcagg**

A portion of the 16S rDNA from *Pseudomonas syringae pv. tomato* DC3000 (3) was amplified with the forward (1) and reverse (2) primers below:

>1___GI-16S_forward_for_16S_G-46647
ACATTCAGATGAAGGCTCAGG**agagtttgatcctggctcag**
>16S_ITSo_reverse_for_16S_G-46648
GTGGTAATGATCCTTCCG**taccttgttacgactt**
>PtoDC3000_16S_27F-1492R
**agagtttgatcctggctcag**ATTGAACGCTGGCGGCAGGCCTAACACATGCAAGTCGAGCGGCAGCACGGGTACTTGTACCTGGTGGCGAGCGGC
GGACGGGTGAGTAATGCCTAGGAATCTGCCTGGTAGTGGGGGATAACGCTCGGAAACGGACGCTAATACCGCATACGTCCTACGGGAGAAAGCA
GGGGACCTTCGGGCCTTGCGCTATCAGATGAGCCTAGGTCGGATTAGCTAGTTGGTGAGGTAATGGCTCACCAAGGCGACGATCCGTAACTGGT
CTGAGAGGATGATCAGTCACACTGGAACTGAGACACGGTCCAGACTCCTACGGGAGGCAGCAGTGGGGAATATTGGACAATGGGCGAAAGCCTG
ATCCAGCCATGCCGCGTGTGTGAAGAAGGTCTTCGGATTGTAAAGCACTTTAAGTTGGGAGGAAGGGCAGTTACCTAATACGTATCTGTTTTGA
CGTTACCGACAGAATAAGCACCGGCTAACTCTGTGCCAGCAGCCGCGGTAATACAGAGGGTGCAAGCGTTAATCGGAATTACTGGGCGTAAAGC
GCGCGTAGGTGGTTTGTTAAGTTGAATGTGAAATCCCCGGGCTCAACCTGGGAACTGCATCCAAAACTGGCAAGCTAGAGTATGGTAGAGGGTG
GTGGAATTTCCTGTGTAGCGGTGAAATGCGTAGATATAGGAAGGAACACCAGTGGCGAAGGCGACCACCTGGACTGATACTGACACTGAGGTGC
GAAAGCGTGGGGAGCAAACAGGATTAGATACCCTGGTAGTCCACGCCGTAAACGATGTCAACTAGCCGTTGGGAGCCTTGAGCTCTTAGTGGCG
CAGCTAACGCATTAAGTTGACCGCCTGGGGAGTACGGCCGCAAGGTTAAAACTCAAATGAATTGACGGGGGCCCGCACAAGCGGTGGAGCATGT
GGTTTAATTCGAAGCAACGCGAAGAACCTTACCAGGCCTTGACATCCAATGAATCCTTTAGAGATAGAGGAGTGCCTTCGGGAGCATTGAGACA
GGTGCTGCATGGCTGTCGTCAGCTCGTGTCGTGAGATGTTGGGTTAAGTCCCGTAACGAGCGCAACCCTTGTCCTTAGTTACCAGCACGTTAAG
GTGGGCACTCTAAGGAGACTGCCGGTGACAAACCGGAGGAAGGTGGGGATGACGTCAAGTCATCATGGCCCTTACGGCCTGGGCTACACACGTG
CTACAATGGTCGGTACAGAGGGTTGCCAAGCCGCGAGGTGGAGCTAATCTCACAAAACCGATCGTAGTCCGGATCGCAGTCTGCAACTCGACTG
CGTGAAGTCGGAATCGCTAGTAATCGCGAATCAGAATGTCGCGGTGAATACGTTCCCGGGCCTTGTACACACCGCCCGTCACACCATGGGAGTG
GGTTGCACCAGAAGTAGCTAGTCTAACCTTCGGGGGGACGGTTACCACGGTGTGATTCATGACTGGGGTG**aagtcgtaacaaggta**

The ITS region of *Hyaloperonospora arabidopsidis* (3) was amplified with the forward (1) and reverse (2) primers below:

>1___16S-ITSo_forward_for_ITSo_G-46649
AAGTCGTAACAAGGTA**cggaaggatcattaccac**
>2___ITSo_reverse_for_ITSo_G-46650
GTGGTGGTGGTGGTGCTCGA**gagcctagacatccactgctg**
>3___ITS1_*H. arabidopsidis*
**cggaaggatcattacca**CACCTAAAAAACTTTCCACGTGAACCGTTTCAACCCAATAGTTGGGGGTCTTATTTGGCGGCGGCTGCTGGCTTAATT
GTTGGCGGCTGCTGCTGAGTGAGCCCTATCAAAAAAAAGGCGAACGTTTGGGCTTCGGCCTGATTTAGTAGTCTTTTTTTCTTTTAAACCCCTT
CCTTAATACTGAATATACTGTGGGGACGAAAGTCTCTGCTTTTAACTAGATAGCAACTTT**cagcagtggatgtctaggct**

The four amplicons were gel purified and combined by overlap extension PCR. First, the ITS amplicon was joined with the 16S rDNA amplicon, and the *GIGANTEA* amplicon was joined with the ITSo amplicon. These two fragments were gel purified and combined by overlap extension PCR to make the final construct below, which was cloned into pGEM®-T Easy (Promega, Madison, WI, USA).

>Synthetic_equimolar_templates
**agaaggagatataccatggcttggtcatttagaggaagtaa**AAGTCGTAACAAGGTTTCCGTAGGTGAACCTGCGGAAGGATCATTATTGAATTA
TGTTTTCTAGATGGGTTGTAGCTGGCTCTTCGGAGTATGTGCACGCCTGTCTGGACTTCATTTTCATCCACCTGTGCACCTTTTGTAGTCTTTT
TCAGGTATTGGAGGAAGTGGTCAGCCTATCAGCTCTTTGCTGGATGTAAGGACTTGCAGTGTGAAAACAGTGCTGTCCTTTACCTTGGCCATGG
AATCTTTTTCCTGTTAGAGTCTATGTTATTCATTATACTCTTAGAATGTCATTGAATGTCTTTACATGGGCTATGCCTATGAAAATTATTATAC
AACTTTCAGCAACGGATCTCTTGGCTCTC**gcatcgatgaagaacgcagctattgaggcgaatcaacctgt**ATCTAACAATCAAACTGCTAACCGT
AAAAGTAGGAATGTCAAGGGACAGGGACCTGTGGCAGCATTTGATTCATACGTTCTTGCTGCTGTTTGTGCTCTTGCCTGTGAGGTTCAGCTGT
ATCCTATGATCTCTGGTCGGGGGAACTTTTCCAATTCTGCCGTGGCTGGAACTATTACAAAGCCTGTAAAGATAAATGGGTCATCTAAAGAGTA
TGGAGCTGGGATTGACTCGGCAATTAGTCATACGCGCCGAATTTTGGCAATCCTAGAGGCACTCTTTTCATTAAAACCATCTTCTGTGGGGACT
CCATGGAGTTACAGTTCTAGTGAGATAGTTGCTGCGGCCATGGTTGCAGCTCATATTTCCGAACTGTTCAGACGTTCAAAGGCCTTGACGCATG
CATTGTCTGGGTTGATGAGATGTAAGTGGGATAAGGAAATTCATAAAAGAGCATCATCATTATATAACCTCATAGATGTTCACAGCAAAGTTGT
TGCCTCCATTGTTGACAAAGCTGAACCCTTGGAAGCCTACCTTAAGAATACACCGGTTCAGAAGGATTCTGTGACCTGTTTAAACTGGAAACAA
GAGAACACATGTGCAAGCACCACATGCTTTGATACAGCGGTGACATCCGCCTCAAGGACTGAAATGAATCCAAGAGGAAACCATAAGTATGCTA
G**acattcagatgaaggctcaggagagtttgatcctggctcag**ATTGAACGCTGGCGGCAGGCCTAACACATGCAAGTCGAGCGGCAGCACGGGTA
CTTGTACCTGGTGGCGAGCGGCGGACGGGTGAGTAATGCCTAGGAATCTGCCTGGTAGTGGGGGATAACGCTCGGAAACGGACGCTAATACCGC
ATACGTCCTACGGGAGAAAGCAGGGGACCTTCGGGCCTTGCGCTATCAGATGAGCCTAGGTCGGATTAGCTAGTTGGTGAGGTAATGGCTCACC
AAGGCGACGATCCGTAACTGGTCTGAGAGGATGATCAGTCACACTGGAACTGAGACACGGTCCAGACTCCTACGGGAGGCAGCAGTGGGGAATA
TTGGACAATGGGCGAAAGCCTGATCCAGCCATGCCGCGTGTGTGAAGAAGGTCTTCGGATTGTAAAGCACTTTAAGTTGGGAGGAAGGGCAGTT
ACCTAATACGTATCTGTTTTGACGTTACCGACAGAATAAGCACCGGCTAACTCTGTGCCAGCAGCCGCGGTAATACAGAGGGTGCAAGCGTTAA
TCGGAATTACTGGGCGTAAAGCGCGCGTAGGTGGTTTGTTAAGTTGAATGTGAAATCCCCGGGCTCAACCTGGGAACTGCATCCAAAACTGGCA
AGCTAGAGTATGGTAGAGGGTGGTGGAATTTCCTGTGTAGCGGTGAAATGCGTAGATATAGGAAGGAACACCAGTGGCGAAGGCGACCACCTGG
ACTGATACTGACACTGAGGTGCGAAAGCGTGGGGAGCAAACAGGATTAGATACCCTGGTAGTCCACGCCGTAAACGATGTCAACTAGCCGTTGG
GAGCCTTGAGCTCTTAGTGGCGCAGCTAACGCATTAAGTTGACCGCCTGGGGAGTACGGCCGCAAGGTTAAAACTCAAATGAATTGACGGGGGC
CCGCACAAGCGGTGGAGCATGTGGTTTAATTCGAAGCAACGCGAAGAACCTTACCAGGCCTTGACATCCAATGAATCCTTTAGAGATAGAGGAG
TGCCTTCGGGAGCATTGAGACAGGTGCTGCATGGCTGTCGTCAGCTCGTGTCGTGAGATGTTGGGTTAAGTCCCGTAACGAGCGCAACCCTTGT
CCTTAGTTACCAGCACGTTAAGGTGGGCACTCTAAGGAGACTGCCGGTGACAAACCGGAGGAAGGTGGGGATGACGTCAAGTCATCATGGCCCT
TACGGCCTGGGCTACACACGTGCTACAATGGTCGGTACAGAGGGTTGCCAAGCCGCGAGGTGGAGCTAATCTCACAAAACCGATCGTAGTCCGG
ATCGCAGTCTGCAACTCGACTGCGTGAAGTCGGAATCGCTAGTAATCGCGAATCAGAATGTCGCGGTGAATACGTTCCCGGGCCTTGTACACAC
CGCCCGTCACACCATGGGAGTGGGTTGCACCAGAAGTAGCTAGTCTAACCTTCGGGGGGACGGTTACCACGGTGTGATTCATGACTGGGGTG**aa**
**gtcgtaacaaggtacggaaggatcattaccac**ACCTAAAAAACTTTCCACGTGAACCGTTTCAACCCAATAGTTGGGGGTCTTATTTGGCGGCGG
CTGCTGGCTTAATTGTTGGCGGCTGCTGCTGAGTGAGCCCTATCAAAAAAAAGGCGAACGTTTGGGCTTCGGCCTGATTTAGTAGTCTTTTTTT
CTTTTAAACCCCTTCCTTAATACTGAATATACTGTGGGGACGAAAGTCTCTGCTTTTAACTAGATAGCAACTTT**cagcagtggatgtctaggctt**
**cgagcaccaccaccaccac**
